# High-confidence structural predictions of extrachromosomal DNA with ecDNAInspector

**DOI:** 10.64898/2025.12.01.691649

**Authors:** Sophia Pribus, Yanding Zhao, Zhicheng Ma, Clemens Weiss, Aziz Khan, Kathleen Houlahan, Christina Curtis

**Affiliations:** Stanford Cancer Institute, Stanford School of Medicine, Stanford, CA 94305; Department of Medicine, Stanford School of Medicine, Stanford, CA 94305; Department of Genetics, Stanford School of Medicine, Stanford, CA 94305; Department of Computational Biology, Mohamed bin Zayed University of Artificial Intelligence (MBZUAI), Abu Dhabi, UAE; McMaster University, Hamilton, ON, Canada; Department of Biomedical Data Science, Stanford University, Stanford, CA 943056; Chan Zuckerberg Biohub, San Francisco, CA, USA

## Abstract

Extrachromosomal DNA (ecDNA) are circularized genomic elements that reside outside canonical chromosomes. ecDNA amplify oncogene copy number, enhance chromatin accessibility, and act as mobile enhancers through cis- and trans-regulatory interactions, collectively boosting oncogene expression. ecDNA has been implicated in tumor progression, intratumoral heterogeneity, and poor patient prognosis. Despite various lines of evidence that ecDNA promotes aggressive disease, the mechanisms and selective pressures leading to ecDNA formation and propagation remain poorly understood as are their structures. While several computational tools have been developed to infer ecDNA presence or absence from short read sequencing data, accurate identification of large or complex ecDNA structures remains challenging. Here we introduce ecDNAInspector, a novel computational framework to systematically assess the confidence of ecDNA predictions from existing inference tools. Leveraging abundant short-read whole genome sequencing (WGS) data from population-scale cohorts, we demonstrate that ecDNAInspector accurately identifies high-confidence ecDNA calls, improving interpretability and facilitating the association with clinical features. As an illustrative example, applied to a cohort of 250 breast cancers, ecDNAInspector identifies associations between ecDNA structure and molecular subgroups of disease. These findings are supported by orthogonal omic data and experimental characterization of ecDNA captured in representative cell lines. ecDNAInspector provides a scalable, data-driven approach to characterize ecDNA structure, enabling integrative studies of the clinical and biological impact of this non-mendelian mode of oncogene amplification and inheritance.

## Introduction

Extrachromosomal DNA (ecDNA) are one- to three- megabase pair circularized genomic regions found outside canonical chromosomes and are associated with shortened patient survival across cancer types^1,2^. ecDNA promotes tumorigenesis by reintegrating into the genome to form somatic rearrangements and increases tumor heterogeneity through abnormal inheritance patterns due to the absence of centromeres^1–3^. Importantly, ecDNA promote oncogene expression by increasing oncogene copy number and chromatin accessibility^2,4,5^ and act as mobile enhancers, employing cis- and trans-regulatory circuits^4,6,7^. Despite various lines of evidence that ecDNA promotes aggressive disease and may be acquired prior to clonal expansion^8–10^, the exact mechanisms and selection pressures leading to ecDNA formation and propagation remain poorly understood. To enable the detailed exploration of ecDNA function and evolution, there is a critical need to elucidate ecDNA structure, particularly in clinico-genomic cohorts where ecDNA features can be associated with disease trajectories and patient outcomes.

Existing tools to identify and predict ecDNA structures are limited to low throughput, high sensitivity assays or high throughput, low sensitivity probability-based inference methods using sequencing data. For example, fluorescence in situ hybridization (FISH) staining for specific genes in cells enriched at metaphase can confirm the presence of those genes on ecDNA, which will be visually distinct from linear chromosomes. Though considered the gold standard in ecDNA detection, this approach depends on prior knowledge and does not fully resolve the sequence structure of the ecDNA. Further, the limited throughput of these methods make them impractical for exploratory, large-scale studies^11,12^. In contrast, inference-based tools, such as AmpliconArchitect^13^ and JaBbA^14^, produce predictions of ecDNA “cycles” based on variations in sequencing coverage and discordant bulk DNA sequencing reads^11^. Each cycle prediction is represented as a series of genomic segments and the paired “breakends” between these segments, forming a single circular ecDNA cycle structure.

Sequencing-based methods have the benefit of being able to leverage a wealth of sequencing data from large cancer genomics cohorts to assess variability in ecDNA presence and structure across individuals and link this variability with clinical annotations. However, in the absence of ground truth data, it is difficult to benchmark ecDNA detection accuracy and sensitivity. Short-read whole genome sequencing (WGS) has reduced accuracy when ecDNA are larger and more complex, hindering the identification of complicated rearrangements and potentially biasing the patterns of structural conservation that are reported^11^. New methods such as CoRAL^15^ and CIDER-Seq^16^ which use long-read sequencing have shown promising initial results in recovering ecDNA in cell lines^4^ but long-read data remains sparse in patient cohorts. Alternative hybrid approaches enable the experimental enrichment of circular structures from nucleic acids, and have been deployed on cell lines and limited patient samples, but do not necessarily improve structural predictions ^4,12^. Given the wealth of patient genomic data from large clinical cohorts such as The Cancer Genome Atlas (TCGA)^17^, the International Cancer Genome Consortium (ICGC)^18^, Hartwig^19^ and others, computational methods to improve the identification and interpretation of complex ecDNA structures from short-read WGS data are warranted.

Here we describe ecDNAInspector, a novel tool designed to assess the confidence of ecDNA predictions produced by existing ecDNA inference tools. ecDNAInspector leverages available short-read WGS data and probability-based methods while improving confidence in computational predictions of ecDNA structures. To evaluate the utility of ecDNAInspector, we applied it to a breast cancer data set, composed of 231 patients in which intersubtype variations of ecDNA frequency, but not structural composition, had previously been characterized^1,8^. ecDNAInspector identifies associations between ecDNA structure and molecular subgroups of disease, findings that are supported by orthogonal omic data and experimental characterization in representative cell lines. ecDNAInspector provides a systematic method for ecDNA structural validation and analysis, enabling higher-confidence utilization of sequencing data-based predictions in studies of ecDNA.

## Results

### Overview of ecDNAInspector

ecDNAInspector provides a systematic method for quality controlled ecDNA inference from WGS, flagging low quality structures for further investigation and selecting high quality structures for downstream analysis. ecDNAInspector takes as input user-provided predictions of ecDNA cycles, defined as the DNA segments and paired breakends that compose putative ecDNA. These predicted cycles are produced by computational tools (e.g. AmpliconArchitect^13^, JaBbA^14^) that infer ecDNA primary structure (i.e., sequence structure) from whole genome sequencing data (**Fig. 1A-1**). ecDNAInspector flags any cycles with suspicious structural qualities, including cycles with extreme size (flagged with the Extreme Cycle Size Boolean [ESB]) and breakends in regions of the genome with low mappability, e.g. high repeat regions, unmappable regions, blacklisted regions, etc. (flagged with the Mapping Error Boolean [MEB]; **Fig. 1A-2**). ecDNAInspector also leverages orthogonal data (e.g. structural variant calls) to support predicted breakpoints in each cycle. To do this, ecDNAInspector compares predicted paired breakends with user-provided structural variant (SV) calls to calculate a True Positive Rate (TPR), False Positive Rate (FPR), and putative False Negative Rate (pFNR) for each cycle (**Fig. 1A-2**). To determine the quality of each cycle, ecDNAInspector performs unsupervised clustering on these quality metrics, from which the user can either choose a cluster of “high confidence” cycles or proceed with an additional optional filtering to select a group of high confidence cycles (which may include “rescued” cycles with desirable features from non-optimal clusters) (**Fig. 1A-3**). These high confidence cycles may then be used for downstream analysis, including built-in ecDNAInspector analysis modules. Thus, ecDNAInspector enables high-confidence utilization of sequencing data predictions in studies of ecDNA structure.

**Figure 1.**
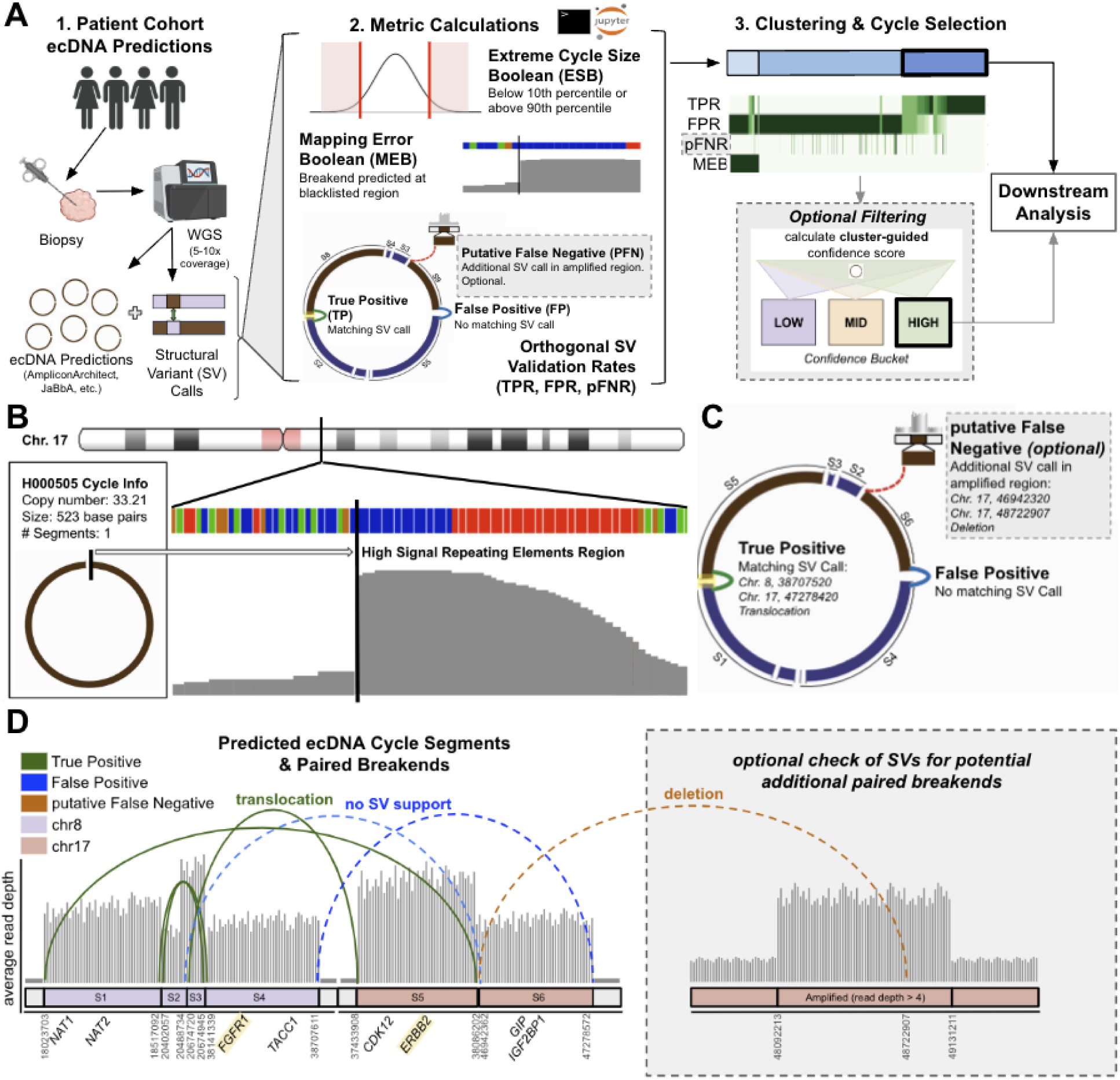
ecDNAInspector uses structural metrics and external validation to nominate low, medium, and high confidence ecDNA cycle predictions. **A.** Schematic shows steps of ecDNAInspector tool. ecDNAInspector takes as input a set of ecDNA cycle predictions from a user-selected prediction tool (1). ecDNAInspector then calculates structure-based metrics for each cycle prediction in the sample set (2) and clusters the cycle predictions based on these metrics (3). The user will select a cluster of high confidence cycle predictions or use an optional filtering step based on clustering results to select a high confidence subset of cycle predictions (3). These high confidence cycle predictions can then be used for downstream analysis leveraging ecDNAInspector analysis modules (4). **B.** Representative schematic showing breakend in a high signal repeating elements region. Cycle predictions with such breakend “mapping errors” are flagged with the Mapping Error Boolean (MEB) metric. **C.** Representative schematic showing breakend validation with orthogonal structural variant (SV) calls. **D.** Representative schematic shows true positive, false positive, and putative false negative paired breakends in the cycle prediction from C). True positive paired breakends are supported by orthogonal SV calls (green solid). False positive paired breakends have no supporting orthogonal SV calls (blue dashed). Putative false negative breakends (orange dahshed) have orthogonal SV support but were not included in the original cycle prediction.

**Figure 2:**
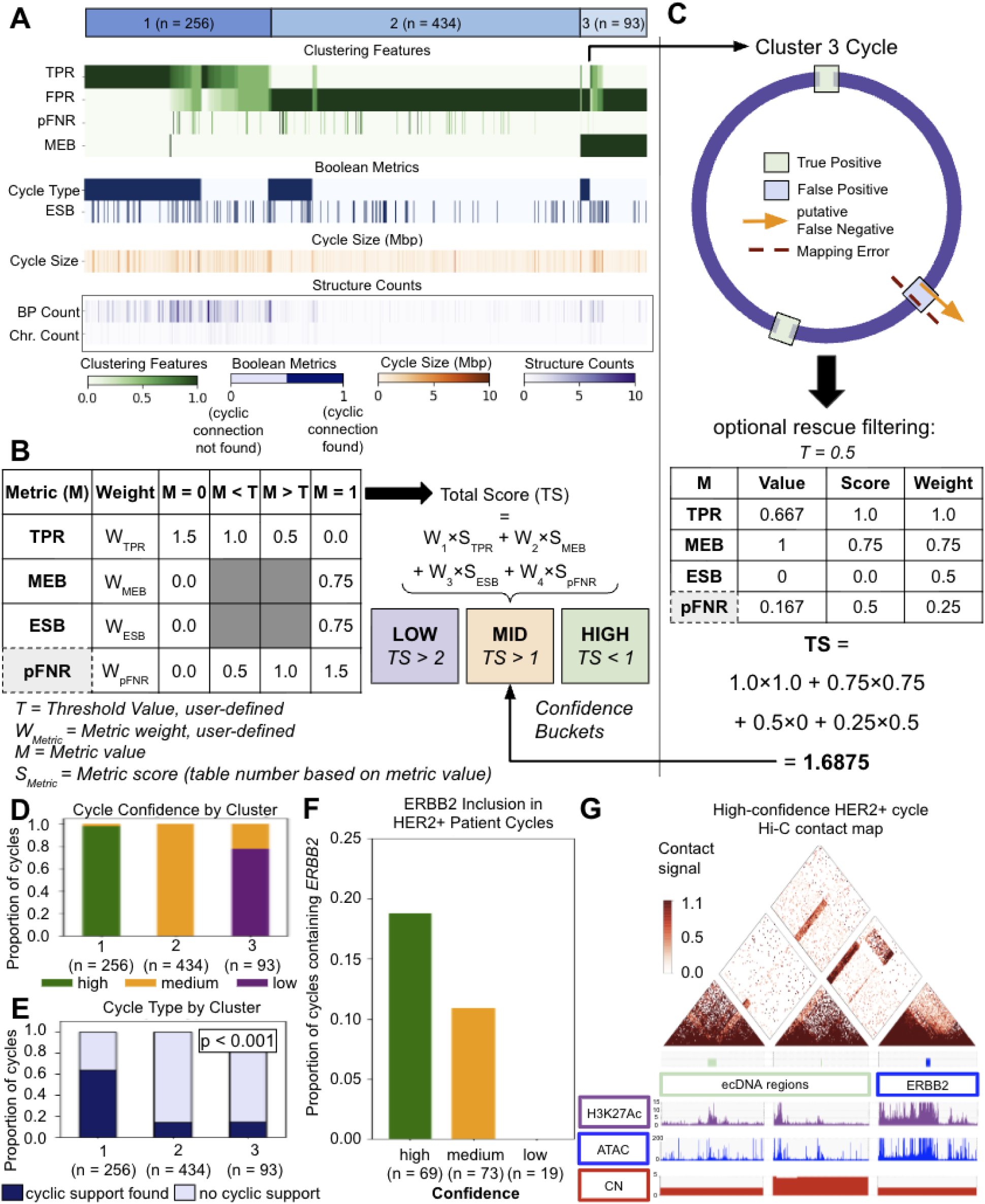
ecDNAInspector nominates high, medium, and low confidence cycle predictions in primary BC cohort. **A**. Heatmap of metrics by cluster for ecDNAInspector clustering on primary BC cohort. Shows distinguishing structural features by cluster. **B**. Schematic of optional post-clustering filtering step. Structural metric values (M) are scored against a user-defined threshold (T). A total score (TS) for each cycle is calculated as a weighted sum of the metric scores. The total score determines ecDNAInspector’s confidence assignment for that cycle. **C**. Example use of filtering step on Cluster 3 cycle prediction, resulting in final assignment as a medium confidence cycle prediction. Table shows metric weight and threshold values. **D**. Post-filtering cycle prediction confidence assignments are plotted by cluster, showing effect of filtering step on re-assigning confidence of cycle predictions. **E**. Pre-filtering cycle cyclic connection support proportions are plotted by cluster, showing significant increase in complete cyclic connection support in cluster 1 (p < 0.001). **F**. *ERBB2* inclusion by cycle prediction confidence for HER2+/IC5 patients. **G**. H3K27ac HiChIP contact matrix (10-kb resolution) in a high confidence HER2+ sample, showing structural proximity of cycle regions.

**Figure 3.**
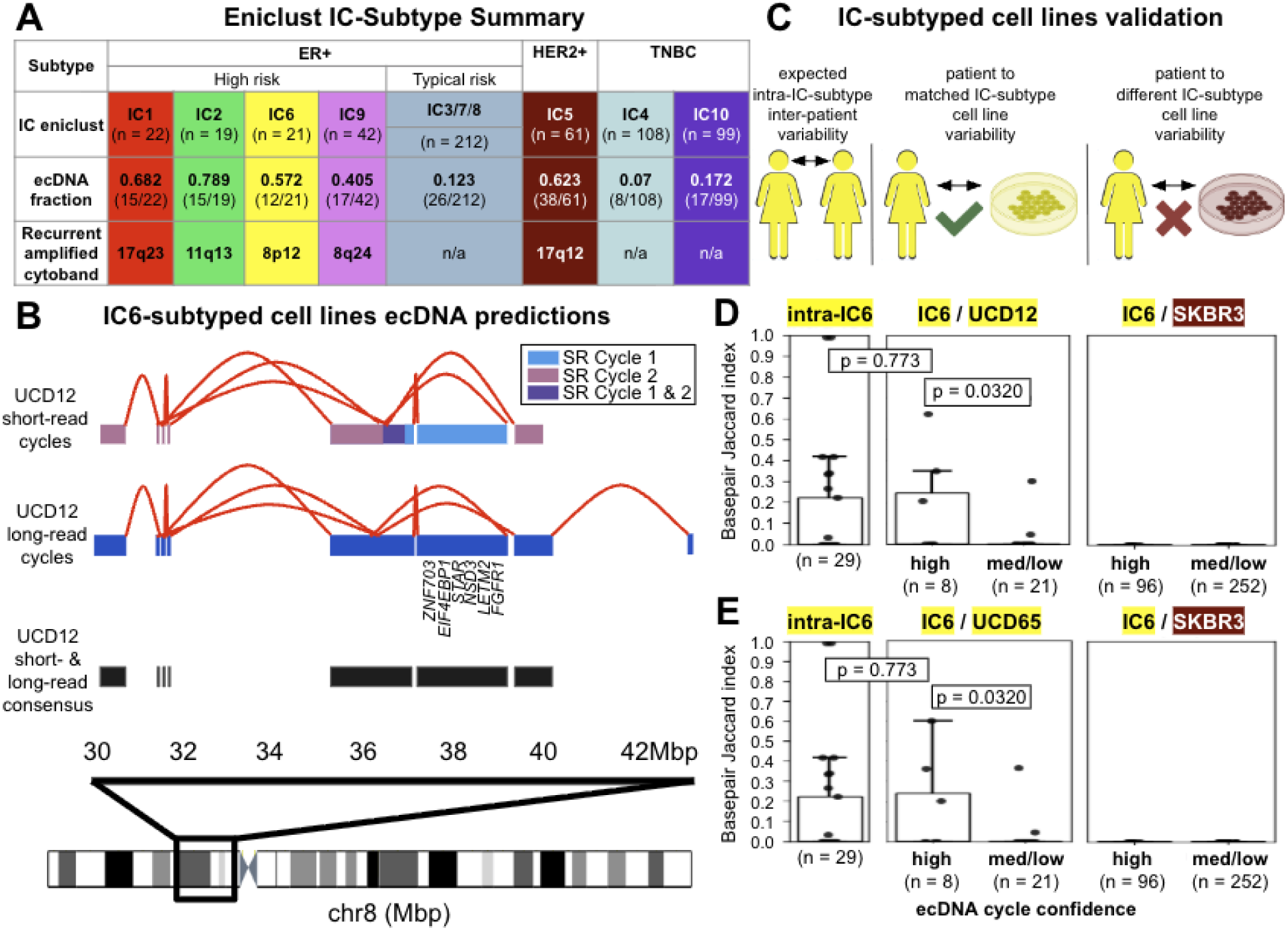
Experimental confirmation of high confidence cycle predictions nominated by ecDNAInspector **A.** Summary of ecDNA prevalence and location by IC ENiClust **B.** A schematic of genomic regions in ecDNA cycle predictions based on short-read and long-read sequencing of the UCD12 cell line. Notable oncogenes are specified. Short-read regions are color-coded by whether they are found in cycle prediction 1, cycle prediction 2, or both cycle predictions from amplicon 2. **C.** A schematic of pairwise cycle comparisons performed to validate ecDNAInspector cycle prediction confidence assignments against expected intra-IC-subtype cycle similarity and cell line-based cycle predictions. **D-E.** Sequence similarity (basepair Jaccard Index) of IC6 high confidence and medium/low confidence cycle predictions to ecDNA regions from the IC6-subtyped UCD12 **(D)** or UCD65 **(E)** cell lines and IC5-subtyped SKBR3 cell line. P-values calculated with Mann-Whitney U-test (two-sided for high confidence/intra-IC6 comparisons, one-sided for high confidence/medium-low confidence comparisons). Boxplots represent median, 0.25 and 0.75 quantiles with whiskers at 1.5x the interquartile range.

### ecDNAInspector identifies 250 high confidence ecDNA cycles from 231 breast tumors

To illustrate the utility of ecDNAInspector to identify high confidence ecDNA predictions, we applied ecDNAInspector to 1,012 ecDNA predictions from a cohort of 231 primary breast cancer patients (primary breast cancer [BC] cohort)^19,20^. Orthogonal SV calls were generated for all samples in the primary BC cohort using the consensus of four SV callers (**methods**). ecDNA cycle predictions were produced using AmpliconArchitect (**methods**). Of note, because the AmpliconArchitect algorithm reconstructs potential ecDNA cycles by identifying unique “walks” through variously connected focally amplified regions, it may produce multiple cycle predictions per sample containing overlapping or identical regions. Prior to any filtering, there was an average of 4.38 “unique” ecDNA predictions per sample. To reduce our cycle set redundancy, we applied ecDNAInspector’s optional intra-sample filter to remove cycles with more than 20% similarity to another cycle in the same sample, keeping the higher quality cycle in the pair (**methods**). This left us with a final set of 783 cycles with matched SV calls for quality control with ecDNAInspector. After this filtering, there was an average of 3.4 “unique” ecDNA predictions per sample.

### Structural and quality metric calculation

ecDNAInspector starts by using the ecDNAInspector_cycle_selection module to parse and calculate structural and quality metrics for each cycle. The structural metrics include cycle segment genomic coordinates, cycle size, cycle chromosome count, cycle breakpoint count, and a list of the cycle’s paired ends (i.e., the segment ends that define a cycle breakpoint). The quality metrics include flags of abnormal size (ESB) and read mapping (MEB) as well as orthogonal SV support (TPR, FPR and pFNR). Cycles with extreme size are defined as any cycle prediction with size below or above a user-defined percentile of the cohort (e.g. <10TH percentile or >90th percentile). ecDNAInspector flags cycles with extreme size (ESB = 1) along with cycles that had any breakends in genomic regions with low mappability (MEB = 1; **Fig. 1B**). ecDNAInspector then leverages orthogonal structural variant calls, defined from a consensus of four SV callers (**methods**), as “ground truth” to classify the paired breakends of each cycle as True Positive or False Positive, and further identified putative False Negative breakends (**Fig. 1C**). True Positive paired breakends are supported by an orthogonal structural variant call, within a user-defined buffer region. These represent predicted ecDNA segment connections with orthogonal support. False Positive paired breakends are not supported by structural variant calls, reducing confidence in the predicted structure of the cycle. If either breakend within a False Positive pair has a different breakend partner identified by structural variant calls in an amplified genomic region, then this is considered a putative False Negative. These represent breakends for which possible ecDNA segment connections are missed by the cycle prediction (**Fig. 1D**).

The distribution of structural metrics can be visualized across the cohort using the ecDNAInspector_analysis module. In the primary BC cohort, prior to quality control filtering, the median cycle size was 0.28 Mbp (standard deviation (s) = 1.541 Mbp; **Supplementary fig. 1A**), and the median breakpoint count per cycle was one (s = 2.06; **Supplementary fig. 1B**). The expected size distribution for ecDNA ranges from 1-3 Mbp. Thus, the unfiltered ecDNA predictions are smaller and less complex than expected. Similarly, the median TPR was 0.0 (s = 0.398; **Supplementary fig. 1C**), suggesting little support from orthogonal SV calls. These metrics indicate a need for quality control filtering.

### Identifying high-confidence cycles

To identify high-confidence cycles, the ecDNAInspector_cycle_selection module performs unsupervised consensus clustering based on the quality metrics. Clustering provides an unbiased grouping of cycle predictions by their structural features, enabling the user to identify high confidence cycle predictions. For our primary BC cohort, ecDNAInspector identified three clusters of cycles (**Fig. 2A, Supplementary fig. 2A-B**). Cluster 1 cycles had medium/high TPR, low/medium FPR, low PFNR, and minimal MEB flags. Cluster 2 cycles had low TPR, high FPR, low/medium PFNR, and minimal MEB flags. Cluster 3 cycles had low TPR, high FPR, low PFNR, and an abundance of MEB flags. These metrics indicated Cluster 1 was enriched for high-confidence cycles. Cluster 1 cycles were significantly larger (p < 0.001, Dunn test; **Supplementary fig. 3A-B**) and had significantly more breakpoints than Cluster 2 and 3 cycles (p < 0.001, Dunn test; **Supplementary fig. 3C**). Cycle size and number of breakpoints alone cannot account for differences in ecDNA prediction quality. That said, the greater size and breakpoint count of Cluster 1 cycles, posited to be enriched for high confidence cycles, are consistent with the observations that ecDNA cycles are large (1-3Mbp)^2^, complex structures with numerous breakpoints.

The ecDNAInspector_cycle_selection module visualizes these structural features of cycles for each cluster (**Fig. 2A**). The user then has an option to select the highest confidence cluster(s) and proceed to downstream analysis, using only the cycles from the selected clusters. Users may also designate the remaining clusters as medium confidence and low confidence clusters. Choosing cycle confidence by cluster alone, with no additional filtering steps, may be suitable when clusters have distinct separation. For example in our primary BC cohort, we assigned Cluster 1, 2 and 3 as high, medium, and low confidence, respectively. Cluster 3 was marked as lower confidence than Cluster 2 because of the enrichment of MEB flags.

Alternatively, the user can use additional optional filtering to select high confidence cycles for downstream analysis. The ecDNAInspector_cycle_selection module enables this optional filtering step with a hierarchical selection process based on a weighted score calculated from the quality metrics (TPR, MEB, ESB, and (optionally) pFNR; **Fig. 2B**). A baseline weight and scoring system for each metric is provided, which was developed by evaluating the stability of various weights and thresholds on our primary BC cohort (see **methods; Supplementary figs. 3D-H**). However, it is highly recommended that the user test multiple weights and thresholds to determine optimal confidence filtering for their personal dataset (i.e., the provided weight and scoring values should not be considered a “default”). ecDNAInspector also offers the user flexibility to prioritize one metric over others or exclude a metric with the weight selection. In our primary BC cohort, applying the ecDNAInspector_cycle_selection module filtering step refined the high confidence cycle set by filtering out medium quality cycles from Cluster 1 while rescuing high quality cycles from Cluster 3 (**Fig. 2C**). Our final confidence set assignments largely followed the original cluster assignments with only 3.32% of cycles reclassified (**Fig. 2D**). In total, we identified 250 high confidence, 460 medium confidence, and 73 low confidence cycle predictions. ecDNAInspector’s additional filtering step enables users to more specifically adjust the confidence assignments of their cycles according to metric priority preferences, if desired.

As noted previously, some inference tools provide all potential ecDNA cycles per sample which may produce cycles containing overlapping or identical regions. To reduce our cycle set redundancy, we applied ecDNAInspector’s optional intra-sample filter to remove highly similar cycles. To ensure this filtering step did not bias confidence assignments, we clustered our original cycle set without intra-sample similarity filtering (**Supplementary fig. 4A**). We found that the intra-sample similar cycle pairs showed significantly greater similarity than all cycle pairs in the full primary BC cohort (p = 0, Rank-Biserial correlation (effect size) = 0.991, two-sided Mann-Whitney U-test; **Supplementary fig. 4B**). The majority of the previously filtered out highly similar intra-sample cycles received the same confidence assignment as their kept partner (**Supplementary figs. 4C-D**). Therefore, ecDNAInspector produces consistent confidence assignments for highly similar cycles.

### High confidence ecDNA cycles are enriched for complete cyclic connections

As an initial validation of ecDNAInspector confidence designations based on clustering alone, we used a confidence flag provided by AmpliconArchitect. For each cycle, AmpliconArchitect reports whether it was able to construct a prediction with complete cyclic connection support (i.e., a sufficient read coverage across all paired breakends in the prediction; a higher confidence prediction) or incomplete cyclic connection support (i.e., one paired breakend lacks read coverage; a lower confidence prediction). In support of our cluster confidence assignments, the proportion of cycles with complete cyclic connection support was significantly higher in Cluster 1 than Clusters 2 and 3 (p < 0.001, Cramer’s V (effect size) = 0.501, Chi-squared test; **Fig. 2E**). Thus, ecDNAInspector’s assignment of cycle confidence by cluster provides a simple unbiased way to refine cycle predictions.

### High confidence ecDNA cycles are enriched for oncogenes

Cycle predictions can be further refined leveraging additional optional filtering via a hierarchical selection process based on quality metrics. As a validation of our confidence designations based on the additional optional filtering, we focused on a subset of ecDNA from HER2-positive (HER2+) patients as ecDNA containing *ERBB2* (encoding HER2) has been well-established; indeed, *ERBB2* is amplified on ecDNA in 26-59% of HER2+ breast cancer patients^1,8^. Thus, as a surrogate of quality, we investigated the proportion or ERBB2 oncogene inclusion in the cycle predictions across confidence groups in HER2+ patients. We found that the proportion of ERBB2 inclusion in cycle predictions from HER2+ patients was highest in the high confidence cycle predictions, reduced in medium confidence cycle predictions, and zero in low confidence cycle predictions (**Fig. 2F**). In breast cancer, which oncogenes are incorporated into ecDNA are highly specific for each breast cancer subtype^8^. An enrichment of oncogene incorporation within high confidence cycle predictions was repeated for other oncogene/genomic subtype pairings (**Supplementary fig. 5A**). This data support the confidence assignments and suggests ecDNAInspector nominates probable ecDNA structures.

### High confidence ecDNA cycles are enriched for three-dimensional contacts

We further performed a Hi-C structural validation of a high confidence cycle prediction and medium confidence cycle prediction from the same HER2+ patient sample. We found that the high confidence cycle showed significant contacts across ecDNA segments from different genomic areas, as well as with the *ERBB2* gene, suggesting mobile enhancer activation of *ERBB2* by ecDNA as previously described in the literature^1^ (**Fig. 2G**). By contrast, the medium confidence cycle showed few cross-segment contacts and poor contact mapping overall (**Supplementary fig. 5B**). These data provide orthogonal support for the confidence assignments further highlighting ecDNAInspectors utility.

### Validation of high confidence ecDNA cycles in breast cancer cell lines

Finally, we validated ecDNAInspector confidence assignments by leveraging breast cancer cell line models of the IC breast subtypes. The IC-subtypes are defined by distinct genomic and transcriptional profiles and are linked to distinct clinical outcomes (**Fig. 3A**)^8^. Importantly, four ER+ IC subtypes (IC1, IC2, IC6 and IC9) have mutational profiles with rearrangement signatures highly concordant with HER2+ tumours and similarly harbor an enrichment of ecDNA. However, instead of amplifying *ERBB2*, this subset of ER+ tumors amplify alternate oncogenes, such as *FGFR1, CCND1* and *MYC* (**Fig. 3A**). Although the breast cancer cell models we used were not derived from the patients in our study, the well-defined patterns of ecDNA across the IC subtypes make them an ideal case study to evaluate ecDNAInspector.

We first assigned each sample in the primary BC cohort to a genomic subtype using the ENiClust algorithm^8^; see **methods**). We observed modest differences in quality metrics and confidence assignments across IC subtypes (**Supplementary figs. 6A-H**). Most notably, ecDNA cycles in IC9 showed the highest pFNR, while IC2 cycles were enriched for ESB flags (**Supplementary figs. 6C&E)**. Overall, IC5 tumors enriched for high confidence cycles, while IC4 and IC10 harbored the most low confidence cycles, as expected given the increased diffuse genome instability in these tumors (**Supplementary fig. 6G**).

Next, we identified consensus ecDNA regions from both short-read and long-read sequencing for two IC6-subtyped cell lines (UCD12 and UCD65; see **methods**). We found strong overlap between the short-read and long-read predictions (**Fig. 3B, Supplementary fig. 7A**), further supporting the value of ecDNA predictions from short-read sequencing. We experimentally confirmed the presence of ecDNA in UCD12 and UCD65 by digestion of linear DNA (**Supplementary fig. 7B**).

To validate the ecDNAInspector confidence assignments, we compared the sequence similarity of ecDNA structures predicted in patients to the experimentally-supported ecDNA from IC-subtype-matched cell lines (**Fig. 3C**). We defined sequence similarity as the ratio of shared to unique bases in the patient vs cell line ecDNA (i.e. basepair Jaccard; **Supplementary fig. 7C**). Compared to medium/low confidence cycle predictions, we anticipated high confidence cycle predictions from IC6 tumors would show higher sequence similarity with ecDNA in the IC6-subtyped cell lines. The IC6 high confidence cycle predictions showed greater sequence similarity with the UCD12 ecDNA regions than the IC6 medium and low confidence cycle predictions (p = 0.032, Rank-Biserial correlation (effect size) = 0.304, one-sided Mann-Whitney U-test; **Fig. 3D, Supplementary fig. 7D**). A similar result was observed for the UCD65 cell line (p = 0.032, Rank-Biserial correlation (effect size) = 0.304, one-sided Mann-Whitney U-test; **Fig. 3E, Supplementary fig. 7E**). Because the UCD12 and UCD65 cell lines are not derived from any of the patients in our primary BC cohort, we would expect some variability in ecDNA cycle sequence structure between the cell lines and the tumor samples. To determine the expected variability, we quantified the intra-IC6 inter-patient sequence similarity (**Fig. 3B**). We found that IC6 patient-UCD12 cell line variability for IC6 high confidence cycle predictions fell well within the expected range of intra-IC6 inter-patient variability (p = 0.773, Rank-Biserial correlation (effect size) = -0.060, two-sided Mann-Whitney U-test; **Fig. 3D, Supplementary fig. 7D**). The same was true for the UD65 cell line (p = 0.773, Rank-Biserial correlation (effect size) = -0.060, two-sided Mann-Whitney U-test; **Fig. 3E, Supplementary fig. 7E**). As a negative control, we compared cycle predictions from IC6 tumors to SKBR3, an HER2+ cell line that harbors ecDNA distinct from IC6 ecDNA (**Fig. 3B**). Neither the IC6 high confidence or IC6 grouped medium and low confidence cycle predictions show any overlap with SKBR3 ecDNA regions (**Figs. 3D-E, Supplementary figs. 7D-E**). These results were not specific to IC6 and were also observed when comparing cycle predictions from IC5/HER2+ tumors against the HER2+ SKBR3 cell line (p = 3.47x10^-5^, Rank-Biserial correlation (effect size) = 0.156, one-sided Mann-Whitney U-test; **Supplementary figs. 7F-G**). The strong subtype specificity of the high confidence cycles indicate ecDNAInspector was able to successfully identify the most probable and supported ecDNA cycles in the IC6 and IC5 subtypes, validating ecDNAInspector’s cycle selection process.

### High confidence cycle subset analysis refines subtype-specific ecDNA structures

Next, we evaluated the impact of confidence filtering on downstream analysis of ecDNA. We first used the ecDNAInspector_analysis module to re-visualize cohort-level metric distributions (see “Structural and quality metric calculation” results section). We found that subsetting to high confidence cycles moved structural metrics into the expected ranges; the cycle size median increased to around 0.91 Mbp (s = 1.804 Mbp; **Supplementary fig. 1D**), much closer to the expected 1-3Mbp range, and the breakpoint count median increased to 3 (s = 2.545; **Supplementary fig. 1E**), a better representation of the expected structural complexity of ecDNA. The TPR median was 1.0 (s = 0.206; **Supplementary fig. 1F**), as expected for high confidence cycles.

Next, we investigated ecDNA differences across IC subtypes in our primary BC cohort. We found no significant difference in cycle size (p_all cycles_ = 0.103, 1²_all cycles_ (effect size) = 0.006, p_high confidence cycles_ = 0.052, 1²_high confidence cycles_ (effect size) = 0.027, Kruskal-Wallis test; **Supplementary Fig. 8A-B**) or cycle breakpoint count (p_all cycles_ = 0.060, 1²_all cycles_ (effect size) = 0.008, p_high confidence cycles_ = 0.025, 1²_high confidence cycles_ (effect size) = 0.035, Kruskal-Wallis test, no significant p-values for high confidence cycle group comparison with Dunn test; **Supplementary Fig. 8C-D**) across the IC subtypes. There was however a significant difference in the gene count per cycle across IC subtypes (p_all cycles_ < 0.001, 1²_all cycles_ (effect size) = 0.027, Kruskal-Wallis test; **Supplementary Fig. 8E**), a trend clarified by examining high confidence cycles only (p < 0.001, 1²_high confidence cycles_ (effect size) = 0.070, Kruskal-Wallis test; **Supplementary Fig. 8F**). We found IC5 cycles had a significantly greater number of genes than IC1 cycles (p = 0.021, Dunn test) and IC10 cycles (p = 0.016, Dunn test). Thus, using ecDNAInspector to select high confidence cycles for downstream analysis enabled clarified exploration of the full primary BC cohort, single cycles of interest within the cohort, and differences between subgroups of the cohort.

### Structural conservation provides insights into ecDNA genesis and function

A refined understanding of ecDNA structure and the conservation of structural features across ecDNA could offer insights into mechanisms of ecDNA development and action. In particular, exploring this conservation in ecDNA from patients across different subgroups (e.g. clinical or genomic subtypes) could improve understanding of subtype-specific disease progression. For example, identifying what common regions or genes are included in ecDNA across subtypes could inform how ecDNA contribute to subtype-specific pathology, while patterns in breakpoint locations could suggest structural vulnerabilities facilitating ecDNA genesis. To address this question, ecDNAInspector quantifies structure conservation by calculating pairwise Jaccard index metrics for various structural features. These include similarity metrics based on shared base pairs (basepair Jaccard index; **Supplementary Fig. 9A**), shared genes (gene Jaccard index; **Fig. 4A**), and shared breakpoints (breakpoint Jaccard index; **Supplementary Fig. 9B**).

**Figure 4.**
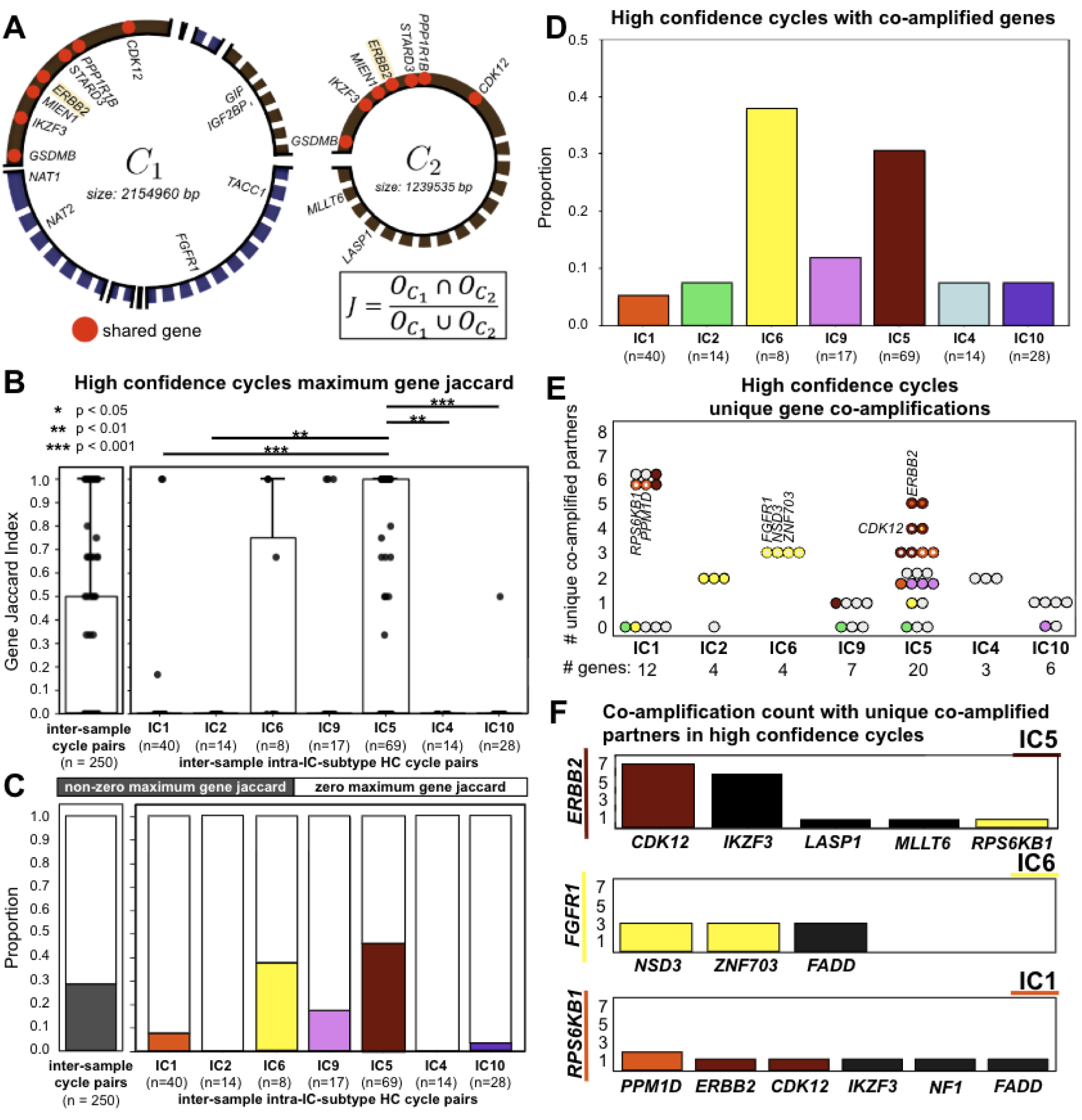
High confidence cycles reveal differences in gene-driven conservation across IC subtypes. **A.** A schematic detailing how pairwise gene Jaccard index metric is calculated. **B.** Comparison of maximum gene Jaccard indices for inter-sample high confidence cycle pairs and intra-subgroup inter-sample high confidence cycle pairs. Boxplots represent median, 0.25 and 0.75 quantiles with whiskers at 1.5x the interquartile range. **C.** Comparison of proportions of non-zero versus zero maximum gene Jaccard indices for inter-sample high confidence cycle pairs and intra-subgroup inter-sample high confidence cycle pairs. **D.** Comparison of proportions of high confidence cycles containing two or more unique genes across IC subtypes. **E.** Count of unique co-amplified genes for each unique gene present on a high confidence cycle across IC subtypes. Each circle represents a unique gene found on a high confidence cycle prediction for that IC subtype. Labeled genes are “IC-specific genes”, defined as genes contained in the recurrently amplified cytoband of an IC subtype. Circles are colored in if 1) a cycle from that IC subtype containing that gene is co-amplified with an IC-specific gene from a different IC subtype, or 2) that gene itself is an IC-specific gene from a different IC subtype. Circles are outlined if a cycle from that IC subtype containing that gene is co-amplified with an IC-specific gene from the same IC subtype. **F.** Count of unique co-amplified gene pairs for selected IC-specific genes for IC5 and IC6 samples. Gene names are underlined and gene bars are colored to indicate their IC-specificity. Each bar represents the number of high confidence cycles from the specified IC subtype in which both the y-axis gene and x-axis gene are present.

### Intra-IC-subtype gene overlap drives ecDNA sequence conservation

We next used ecDNAInspector’s cohort analysis module to evaluate ecDNA structure conservation across breast tumors, considering only high confidence cycles. We found 19.2% of cycles had a maximum basepair Jaccard index equal to zero, indicating no sequence similarity with any other cycle. Amongst the rest, there was a high degree of variability in structure conservation (**Supplementary Fig. 9C**). To better understand the source of this conservation, we explored whether it could be explained by conserved gene inclusion or conserved breakpoints.

The former may indicate conserved ecDNA structure is driven by gene selection pressures, while the latter could suggest this conservation is the result of a mechanism of ecDNA genesis from recurrent fragile sequence positions. We found that basepair Jaccard indices were significantly greater than gene Jaccard indices (p = 1.16x10^-11^, Rank Biserial correlation (effect size) = 0.360, two-sided Mann-Whitney U-test) and breakend Jaccard indices (p = 6.12x10^-34^, Rank Biserial correlation (effect size) = 0.630, two-sided Mann-Whitney U-test). Gene Jaccard indices were also significantly greater than breakend Jaccard indices (p = 7.64x10^-4^, Rank Biserial correlation (effect size) = 0.134, two-sided Mann-Whitney U-test) (**Supplementary Fig. 9C**). These data point to oncogene-driven selection pressure over sequence fragility as a driving factor in ecDNA structure. However, the high basepair conservation suggests non-genic regions, which may include regulatory elements, may also be conserved^7^.

Given the well-defined patterns of ecDNA regions and oncogene inclusion across IC-subtypes, we wanted to confirm whether this conservation was driven by intra-IC-subtype comparisons. Comparing cycles from the same subtype, we found similar patterns in basepair, gene, and breakend Jaccard (**Supplementary 9C-D**), indicating intra-IC-subtype comparisons drive cycle similarity. Across IC subtypes, we found significant differences in basepair conservation (p = 3.42e-9, 1² (effect size) = 0.184, Kruskal-Wallis test; p = 2.08e-7, Cramer’s V (effect size) = 0.469, Chi-squared test; **Supplementary Fig. 9E-F**) and oncogene conservation (as defined by COSMIC; p < 0.001, 1² (effect size) = 0.153, Kruskal-Wallis test; **Figure 4B**), but not breakend conservation (p = 0.241, 1² (effect size) = 0.008, Kruskal-Wallis test; p = 0.377, Cramer’s V (effect size) = 0.202, Chi-squared test; **Supplementary fig. 9G-H**). In particular, the IC2 and IC5 subgroups show strong basepair conservation in comparison to the IC1, IC9, IC4, and IC10 subgroups (Dunn test; see code for exact Dunn test p-values). Interestingly, oncogene overlap appeared to explain the conservation trend in IC5 cycles, but not IC2 (**Fig. 4C**). Breakend overlap was also not conserved in the IC2 cycles (**Supplementary figs. 9G-H**). Thus, while IC5 ecDNA conservation may be explained by intra-IC-subtype oncogene recurrence, IC2 ecDNA conservation cannot be explained by either gene or breakend recurrence. Thus, ecDNAInspector can identify structural differences that hint at subgroup-specific mechanisms of ecDNA generation and action.

While ecDNA structure conservation was highest amongst cycles from the same subtype, we also observed structure conservation across subtypes (**Supplementary fig. 9C**). To explore which genes drove both intra- and inter-IC-subtype gene conservation across IC subtypes, we identified oncogene co-amplification (defined as two or more unique genes present on the same ecDNA cycle prediction) amongst the high confidence cycles. We found that one third of IC5 high confidence cycles and over one third of IC6 high confidence cycles had gene co-amplification, with more limited co-amplifications in IC1, IC2, IC9, IC4, and IC10 (**Fig. 4D**). We found frequent co-amplifications of IC1 cytoband genes and IC5 cytoband genes in IC5 ecDNA (**Fig. 4E)**. IC6 co-amplifications occurred amongst IC6 cytoband genes. Across all subtypes, cycles that contained *CCND1*, an IC2 cytoband gene, were significantly depleted for co-amplifications compared to cycles containing other IC cytoband genes (p = 0.006, OR=0.05, Fisher’s Exact Test). Overall, we identified the most frequent unique co-amplifications in the IC1, IC5, and IC6 subtypes (**Fig. 4F**). We found that IC5 ecDNA showed frequent co-amplification of *ERBB2* and another IC5 gene, *CDK12*, and less frequent co-amplifications of *ERBB2* with an IC1 gene, *RPS6KB1* (**Fig. 4E-F**). IC6 ecDNA show moderate co-amplifications between *FGFR1* and other IC6 genes, *NSD3* and *ZNF703* (**Fig. 4E-F**). IC1 ecDNA show infrequent co-amplifications between *RPS6KB1* and another IC1 gene, *PPM1D*, and IC5 genes, *ERBB2* and *CDK12* (**Fig. 4E-F**). Finally, given our high frequency of *ERBB2* co-amplifications, we wanted to check beyond oncogenes to see if this extended to immune genes previously identified as frequently included in ecDNA. We identified one high confidence cycle with co-amplification of *ERBB2* and immune genes *GIT1, CCL1,* and *CCL13* (out of 21 total high-confidence cycles with *ERBB2*). Thus, by using ecDNAInspector for cycle selection and downstream analysis, we revealed that ecDNA structural conservation is primarily driven by IC-specific gene inclusion, with distinct co-amplifications driving inter-subtype conservation. In particular, we identified that *ERBB2* frequently co-amplified with other cancer genes across multiple subtypes, and in one case, with important immune genes.

## Discussion

The presence of ecDNA is emerging as a hallmark of tumors; however, the repertoire of mechanisms by which they promote tumorigenesis remains to be seen. Further, questions remain regarding ecDNA genesis and maintenance during tumor growth. Given the tight link between structure and function in biology, precise measurements of ecDNA structure across tumors and cancer types will help provide answers to these questions. Inference-based methods are able to leverage the wealth of available tumor sequencing data and provide ecDNA predictions concordant with more experimentally laborious techniques optimized for circular DNA detection, *e.g.* CIRCLE-seq, CRISPR-CATCH^5,21^. Indeed, here we demonstrate strong concordance between ecDNA inference from short and long read sequencing on two different cell lines that was further confirmed by experimentally depleting linear DNA. These data indicate that inference based tools can identify the presence of ecDNA, however, their comprehensive outputs can also include a large number of small, low complexity predicted cycles that are likely artifacts. ecDNAInspector refines these outputs to identify high confidence cycle predictions for deeper discovery of ecDNA structure and function.

As proof of principle, ecDNAInspector nominated 250 high confidence ecDNA cycles from 231 breast tumors and validated these high confidence cycles by patient-matched orthogonal three-dimensional chromatin conformation data and subtype-matched cell line profiling. Previously, we discovered four subtypes of ER+ breast cancer that were enriched for ecDNA, similar to HER2+ tumors^8,22^. However, instead of focally amplifying *ERBB2*, these ER+ tumors harbor complex amplifications of alternate oncogenes, including *FGFR1, CCND1* and *MYC*. Leveraging ecDNAInspector, we discovered ecDNA cycles within these subtypes show a high degree of structural conversation, driven predominantly by conserved oncogene incorporation. These data suggest oncogene-driven selection pressures rather than sequence fragility drives ecDNA genesis and maintenance. Structure conservation was highest amongst tumors of the same subtype, with IC5 and IC6 ecDNA showing the highest conservation. However, inter-subtype conservation was also observed, largely driven by IC5 ecDNA displaying sequence similarity with non-IC5 ecDNA. Further investigation showed this is due to frequent *ERBB2* co-amplification with other IC-specific oncogenes. Taken together, these data motivate oncogene selection as a driving force in ecDNA genesis and maintenance with the number of oncogenes selected for ranging across subtypes.

Notably, our structural conservation analysis revealed no gene overlaps in the IC2 subtype, an unexpected result given their well-defined genomic amplifications in genic regions. *CCND1*, an IC2-specific oncogene, was not present in any high-confidence IC2 cycles, and the depletion of co-amplifications in non-IC2 cycles containing *CCND1* was suspicious. On further investigation, we found strong *CCND1* conservation in medium confidence IC2 cycles (**Supplementary fig. 5A**). It has been previously shown that the IC2 subtype presents with a lower SV burden^8^; thus it is plausible that without significant orthogonal SV calls with which to support IC2 cycle predictions, IC2 cycles were disproportionately marked to a lower confidence group. Thus, in this primary BC cohort it would be reasonable to analyze both high and medium confidence IC2 cycles for biological discovery.

This example demonstrates that user diligence is necessary for successful ecDNAInspector application. The user is required to carefully investigate cycle confidence assignments as optimal parameter selections are likely cohort-specific. ecDNAInspector provides easy to use and flexible modules that support these selections, including an optimal filtering scheme with flexible weighting to prioritize or deprioritize specific quality metrics. Further, the quality of ecDNAInspector filtering is highly dependent on the quality of input data used for orthogonal validation. Ensuring high confidence structural variant detection will help ensure high confidence ecDNA detection. By following these best practices, ecDNAInspector ensures robust and reliable computational predictions of ecDNA structures, enabling direct connections between ecDNA architecture, tumor phenotype, and clinical outcomes.

## Acknowledgements

This research was supported by the National Cancer Institute through the Metastasis Research Network U54 award (CA261719) and a Breast Cancer Research Foundation Award to C.C. We acknowledge the use of data from the Hartwig Medical Foundation and the TCGA Research Network (https://www.cancer.gov/tcga). This research was made possible through access to data in the National Genomic Research Library, which is managed by GEL (a wholly owned company of the Department of Health and Social Care). The National Genomic Research Library holds data provided by patients and collected by the NHS (National Health Service) as part of their care and data collected as part of their participation in research. The National Genomic Research Library is financed by the National Institute for Health Research and NHS England. The Wellcome Trust, Cancer Research UK and the Medical Research Council have also financed research infrastructure.

## Author contributions

Conceptualization: S.J.P, K.E.H, and C.C. Bioinformatic analysis: S.J.P and Y.Z. Statistical analysis: S.J.P. ecDNAInspector tool conceptualization and development: S.J.P., K.E.H, C.W, A.K. Data processing: A.K. Experimental validation: Z.M. Supervision: K.E.H. and C.C. Writing (original draft): S.J.P and K.E.H. Writing (review and editing): all authors.

## Materials and Correspondence

Correspondence to Kathleen E. Houlahan and Christina Curtis.

## Methods

### Generation of Starter Data

We leveraged whole genome sequencing data for 1,422 paired tumor and normal breast cancer samples^20^ encompassing primary invasive disease. All tumor samples were IC-subtyped using the ENiClust algorithm^22^. These samples are referred to as the primary BC cohort.

### Inferring ecDNA structures from whole genome sequencing

We used AmpliconArchitect^13^ (AA, v1.2_r2) to generate cycle predictions for each of the samples in the primary BC cohort. AA uses short-read whole genome sequencing (WGS) data to infer the architecture of focal amplifications. First, candidate amplifications were detected using CNVKit batch^23^ (v0.9.9) with the following parameters: –diagram –scatter –annotate refFlat.txt – access access-hs37d5-cnvkit.bed –method wgs. We then used the refFlat.txt annotation file from UCSC (http://hgdownload.soe.ucsc.edu/goldenPath/hg19/database/refFlat.txt.gz), and generated the the access file (access-hs37d5-cnvkit.bed) using cnvkit.py access hs37d5.fa -o access-hs37d5-cnvkit.bed. Amplifications with a copy number > 4 were used to seed AA by first trimming segments using seed_trimmer_v2.py within the PrepareAA module setting --minsize 50000. We then ran PrepareAA.py on the trimmed seeds using the following parameters: --cngain 4.0 --cnsize_min 50000 --downsample 10.

### ecDNAInspector Tool Design

ecDNAInspector can be run as a series of Jupyter notebooks or downloaded as a package and run through the command line. Both versions include a series of core modules: cycle metric calculation, cycle clustering, and cycle confidence assignment. Both versions also include an optional intrasample filtering module, and the Jupyter notebook version additionally includes Jaccard calculation and analysis modules. These modules are introduced briefly here and described in further detail below:

1. **Cycle metric calculation (core)**: Takes as input raw cycle prediction files (including at minimum cycle segment, paired end, and copy number information) and outputs a cycle-level data table with all calculated cycle metrics (detailed below). ecDNAInspector converts output data files from ecDNA inference tools into a compatible format and extract structural features to calculate cycle-level metrics. *Included in ecDNAInspector_cycle_selection Jupyter notebook, --run-metric-calc CLI flag*.
2. **Cycle clustering (core)**: Takes as input the cycle-level data table and clusters cycles based on user-specified metrics. The clustering results can then be visualized. *Included in ecDNAInspector_cycle_selection Jupyter notebook, --run-cluster and --visualize_cluster CLI flags*.
3. **Cycle confidence assignment (core)**: Takes as input the Takes as input the cycle-level data table and/or cluster assignments and assigns each cycle to a confidence group. This can be done directly by assigning each cluster to a confidence group or through a hierarchical selection process. *Included in ecDNAInspector_cycle_selection Jupyter notebook, --run-conf-assignment CLI flag*.
4. **Intrasample filtering (optional)**: Users may optionally filter the data to reduce similar cycles from a single sample (intra-sample filtering step). Note, this intra-sample filtering step is more specific for AA ecDNA predictions which output all possible cycles. *Included in ecDNAInspector_cycle_selection Jupyter notebook, --run-intrasample-filter CLI flag*.
5. **Jaccard calculations (Jupyter notebook only):** Takes as input the cycle-level data table from and outputs a new cycle-level Jaccard table calculating the basepair, gene, and breakend Jaccard indices for each pair of cycles in the cohort. This step is optional and designed to provide insights into structural conservation across samples. See “Jaccard calculations module”. *Included in ecDNAInspector_Jaccard_calculations Jupyter notebook*.
6. **Analysis (Jupyter notebook only):** Takes as input the cycle-level data table and (optionally) the cycle-level Jaccard table and provides comparative analyses. Three main analysis sections include: full cohort metric analysis, single cycle versus cohort analysis, and subgroup analysis (requires additional input from users with desired subgroup assignments for all samples). *Included in ecDNAInspector_analysis Jupyter notebook*.

The Jupyter notebooks include: ecDNAInspector_cycle_selection, ecDNAInspector_Jaccard_calculations, ecDNAInspector_analysis, and ecDNAInspector_functions. ecDNAInspector_functions codes calculation and plotting functions used by the remaining three notebooks. The remaining three notebooks should be run in the order defined above. The command line tool run flags include: --run-metric-calc, --run-cluster, --run-conf-assignment, --run-intrasample-filter, and --visualize_cluster. Note that the command line version can be used to calculate the cycle metrics, cluster the cycles, and assign confidence levels to the cycles, and the output can then be used in the ecDNAInspector_analysis Jupyter notebook for further visualizations. For more details on both versions, please see the ecDNAInspector github page [link].

### Cycle metric calculation module

#### Required file inputs

The **cycle metric calculation** module requires the following input files:

● Output data files from ecDNA inference tools for each sample
● An Amplicon information file, where each row includes the sample name and an amplicon number (to designate the different amplicons to be considered for each sample)
● A BEDPE file containing orthogonal structural variant calls for all cohort samples
● A file containing information on genes to be identified in cycles (default provided)
● A file containing information of blacklisted genome regions to be flagged in cycles (default provided)

#### Raw data file conversion

The **cycle metric calculation** module supports conversion of raw ecDNA prediction data files from AmpliconArchitect. A standard cycle data file format is used in ecDNAInspector, so users can convert their own raw files if they use a different prediction tool. The standard cycle data file is a csv including columns for:

● The cycle number (“CycleNum”): a unique numeric identifier for each cycle
● The cycle type (“CycleType”): for AmpliconArchitect input, indicates whether the cycle has complete cyclic support (CycleType = “circular”) or incomplete cyclic support (CycleType = “linear”).
● The cycle information (“SegmentsTuples(chr,start,end)”; may be more than one column): a series of tuples describing each segment in the cycle, including their chromosome, starting basepair, and ending basepair. Note these tuples must be read from left to right to reconstruct the cycle segment connections; that is, for each tuple, the “end” basepair is connected to the “start” basepair of the following tuple.

### General structural feature extraction

The **cycle metric calculation** module uses text file extraction to pull out cycle segment information for all amplicon structure predictions for each sample. It includes only cycle predictions with a copy number greater than a user-provided limit. For our primary BC cohort analysis, we included only cycle predictions with copy number > 4.0. Even within regions predicted to harbour ecDNA, AmpliconArchitect additionally provides information on whether an individual cycle has full cyclic connection support (i.e., a full circular cycle was predicted from the AmpliconArchitect breakpoint graph; denoted as “circular” cycles) or incomplete cyclic connection support (i.e., the end-points of the cycle predicted from the AmpliconArchitect breakpoint graph are connected to undetermined positions or positions outside the amplicon interval set; denoted as “linear” cycles). Note, linear cycles are different from amplicons that are predicted to be linear. If using AmpliconArchitect cycle files as input, the user can additionally specify whether to include only cyclic cycles, only linear cycles, or both cyclic and linear cycles. For our primary BC cohort analysis, we included both cyclic and linear cycles.

### Mapping Error Boolean determination

To identify regions with potential mappability issues, ecDNAInspector uses a user-provided genome blacklist and identifies cycle breakends within a user-provided buffer region of problematic regions. Problematic regions include regions with low mappability, high signal, high repeats, or other anomalies resulting in consistent poor quality mapping results. Mapping errors are stored as a Boolean for each cycle; if a cycle has a single mapping error at one of its breakends, that cycle is marked Mapping Error Boolean (MEB)-positive. For our primary BC cohort analysis, we used the hg19 Genome Blacklist (https://github.com/Boyle-Lab/Blacklist/tree/master/lists) and a buffer region of 100 base pairs.

### Cycle gene identification

To identify genes in cycles, ecDNAInspector uses a user-provided set of gene locations and identifies genes that have above a user-provided proportion of their length included in each cycle. For our primary BC cohort analysis, we used the hg19 v98 Cosmic Cancer Gene Census (https://cancer.sanger.ac.uk/cosmic/download/cosmic/v98/cancergenecensus) and included only genes where 100% of the gene length was on cycles.

### External structural variant call validation

To quantify the degree to which ecDNA breakpoints are supported by orthogonal SVs, ecDNAInspector calculates three metrics on a per-cycle basis, visualizations of which are found in **figures 1C, D**:

1. The **True Positive Rate (TPR)** is calculated by finding the number of breakpoints in the cycle that are supported by SV calls (i.e., there is an SV call that connects the paired ends making up the ecDNA breakpoint) and dividing by the total number of breakpoints in the cycle.
2. The **False Positive Rate (FPR)** is calculated by finding the number of breakpoints in the cycle that are not supported on either end by SV calls and dividing by the total number of breakpoints in the cycle.
3. The **putative False Negative Rate (pFNR)** is calculated by first subsetting to the paired breakends of False Positive breakpoints. For each breakend, if it is supported by an SV and the other breakend of the SV is within a user-provided buffer of an amplified region (i.e., a region with copy number greater than the minimum user-provided copy number for cycle prediction inclusion), then the ecDNA breakend is marked as a Putative False Positive. This is because the connection in the predicted ecDNA structure is not supported by SV. Instead, there is evidence of an additional amplified segment that is missed. The total count of these putative false negative breakends is divided by the total number of breakends in False Positive breakpoints. For our applied analysis, we used a buffer of 100 base pairs for all calculations of pFNR.

Because many structural variant calling algorithms are not precise at base pair resolution, there is an option to exclude any deletion-type paired breakends in the ecDNA cycle prediction from the above analysis. ecDNAInspector will exclude any of these paired breakends with less than a user-provided amount of distance between them. For our primary BC cohort analysis, we used a limit of 2000 base pairs (i.e., paired breakends had to be at least 2000 base pairs apart to be included in calculations of TPR, FPR, and pFNR). SV calls were detected using a consensus approach across 4 independent callers: MANTA^24^ (v1.6.0), DELLY^25^ (v0.9.1), SvABA^26^ (v1.1.3), and GRIDSS^27^ (v2.13.2). MANTA was run on tumor/normal pairs using default settings. DELLY was run on tumor/normal pairs excluding unmappable regions annotated in the DELLY blocklist using -x hg19.excl. SvABA was run on tumor/normal pairs using default parameters and excluding ENCODE blacklist regions (--blacklist hg19-blacklist-nochr.v2.bed) and annotated with dbSNP (v138) (--dbsnp-vcf dbsnp_138.b37.vcf). Finally, GRIDSS was run on tumor/normal pairs using the following parameters: --blacklist hg19-blacklist-nochr.v2.bed – picardoptions VALIDATION_STRINGENCY=LENIENT. Somatic SVs called by GRIDSS were filtered with the gridss_somatic_filter function using default parameters. SV calls were kept if they were identified by GRIDSS and at least one other SV caller (MANTA, DELLY, or SvABA).

### Cycle clustering module

#### Unsupervised clustering with structural validation metrics

The **cycle clustering** module uses the a consensus clustering implementation package (github.com/ZigaSajovic/Consensus_Clustering/blob/master/consensusClustering.py) to cluster cycles with KMeans clustering based on structural validation metrics (TPR, FPR, pFNR, and MEB). This package is a Python implementation of a previously published consensus clustering algorithm’s methods to find the ideal K using multiple metrics^28^. Specifically, K is selected by finding the K with greatest consensus over multiple runs of the clustering algorithm with random restart to account for sensitivity to the starting conditions of each run. Users specify the minimum and maximum number of clusters to attempt (default minimum = 3, no default maximum), the number of cluster resamples (default = 1000), the proportion of samples to resample (default = 0.8), and the number of Kmeans runs to perform with different centroid seeds (default = 10). The K-means clustering package is imported from sklearn (v1.6.1). We first tested the clustering using a minimum of 2 and maximum of 8 clusters, with all other settings at the default. For each value of K the consensus clustering algorithm computed a N by N consensus matrix storing the fraction of iterations that any two samples clustered together, where N was the total number of samples. This matrix was then flattened and its CDF calculated. We made an elbow plot of the area under the CDF curve (AUC; (**Supplementary fig. 10A**) and a knee plot of the change in AUC (**Supplementary fig. 10B**), and identified the inflection points.

The AUC indicates the stability of the clustering for a specific K, while the change in AUC indicates how much the stability improves at a specific K. These evaluations confirmed K = 3 as the ideal number of clusters, with larger K offering only marginal improvements in clustering stability. We also compared these results with a separate K-means clustering with the selected value of K = 3, finding an Adjusted Rand Index (ARI) of 0.7313.

### Cycle confidence assignments module

#### Cluster-based method

The **cycle confidence assignments** module can assign each cluster of cycles to a cycle confidence group. Users can use the cluster visualization to visually determine clusters with high, medium, and low confidence qualities to determine which clusters belong to which confidence group.

#### Optional filtering selection method

For a more fine-tuned approach, the **cycle confidence assignments** module offers an optional filtering step to make cycle confidence assignments with a hierarchical selection method. Users define a TPR threshold, a pFNR threshold, score sets and weights for TPR, pFNR, MEB, and ESB, and total score thresholds. Default values are provided and are as follows:

● TPR threshold = 5
● pFNR threshold = 5
● TPR score set: [1.5, 1.0, 0.5, 0.0]
● pFNR score set: [0, 0.5, 1.0, 1.5]
● MEB score set: [0, 0.75]
● ESB score set: [0, 0.75]
● Total score thresholds: [1, 2]

The TPR/pFNR threshold and score sets are used to assign an adjusted value to each of TPR, pFNR, MEB, and ESB (one of the “scores” in their respective set, assigned according to the original value). These metric scores are multiplied by their respective weights and added, and the sum compared to the total score thresholds that define high, medium, and low confidence cycles. A visualization and example of this technique is provided in **figures 3B-C**. We used this filtering step on our primary BC cohort analysis, using the default score sets, weights, and thresholds.

### Optional filtering method output stability evaluation

To confirm stability of the filtering method outputs across score set, threshold, and weight metric choices, we tested a range of these metrics in combination (**Supplementary Fig. 2E**) and computed how many high confidence cycle predictions they resulted in. To first eliminate any metric choices that disproportionately led to no high confidence cycles being identified, we determined the count of metric combinations resulting in a zero high confidence cycle count while varying the value of one metric at a time. We defined a range of acceptable proportions of the total metric combination count that could result in zero high confidence cycles; at minimum, 10% of the combinations while holding a given metric constant should result in zero high confidence cycles (so the metric value was not too permissive), but at maximum 20% of the combinations while holding a given metric constant could result in zero high confidence cycles (so the metric value was not too strict). We eliminated potential values for metrics that resulted in their zero high confidence count going outside this range (i.e., low and high scores for the total score threshold, high TPR scores, and high pFNR scores) (**Supplementary Fig. 2F**). We then determined the range of the count of high confidence cycles produced while varying the TPR/pFNR threshold (**Supplementary Fig. 2G**) and the weights (**Supplementary Fig. 2H**) to confirm the distributions of count of high confidence cycles were relatively consistent across different metric values. We finally held all score sets at the medium value and determined the count of high confidence cycles produced by different combinations of the TPR/pFNR threshold and weights, identifying multiple combinations as producing a similar number of high confidence cycles to other combinations, indicating the specific stability of output across metric choices when holding score sets constant at medium value. This led us to select the default filtering metric values as medium score sets, medium TPR/pFNR threshold, and gradient weights.

### Optional intra-sample similarity filtering module

ecDNAInspector’s optional intra-sample similarity filter removes some cycle predictions to reduce highly similar cycle predictions from the same sample. This filter is specifically designed for the output of inference tools that may produce multiple similar cycle predictions per sample (e.g., AmpliconArchitect). This filter uses the basepair Jaccard index metric (see “ecDNAInspector_Jaccard_calculation module”) to identify intra-sample groups of cycle predictions with more than a user-defined threshold of similarity. For each of these groups of cycle predictions within unique samples, only the higher quality cycle prediction (the cycle with greater TPR) is kept for validation and downstream analysis. We applied this filter to our primary BC cohort and used a threshold of 20% similarity (i.e., cycle pairs with basepair Jaccard index > 0.2 were filtered to choose the single higher quality cycle). We confirmed this accurately selected highly similar cycles from the cohort and checked it did not affect the clustering or confidence assignments by reclustering without filtering and comparing the confidence assignments of similar intra-sample cycles (**Supplementary Figs. 3A-D**).

### Jaccard calculation module

### Required file inputs

The **Jaccard calculation** module requires the cycle level data table, including all calculated structural metrics for each input cycle (specifically the cycle regions list and cycle genes list). The cycle level data tables with cluster/confidence assignments or the intra-sample filtered cycle level data table can equivalently be used.

### Jaccard index calculations

ecDNAInspector calculates three main similarity metrics for pairwise comparisons: the base pair Jaccard Index, oncogene Jaccard Index, and breakend Jaccard Index.

1. **The base pair Jaccard Index** is calculated by finding the intersection of all base pairs across two cycle predictions and dividing by the union of all base pairs across two cycle predictions. See **Supplementary fig. 8A.**
2. **The gene Jaccard Index** is calculated by finding the intersection of all genes present across two cycle predictions and dividing by the union of all genes across two cycle predictions. The proportion of the gene overlapping a cycle region must be greater than a user-provided fraction for the gene to be considered present in the cycle. For our analysis, we used a proportion of 1.0 (meaning the entire gene had to be present in the cycle to be considered included). See **Figure 4A**.
3. **The breakend Jaccard Index** is calculated by finding the count of overlapping breakends across two cycle predictions and dividing by the total number of breakends across two cycle predictions. Breakends must be within a user-provided buffer to be considered overlapping. For our analysis, we used a buffer of 100 base pairs. See **Supplementary fig. 8B**.

### Output data file

The **ecDNAInspector_Jaccard_calculation** module outputs a single Jaccard index information file. Each row represents a comparison between two cycles and includes the following information: sample 1 name, sample 1 amplicon, sample 1 cycle number, sample 2 name, sample 2 amplicon, sample 2 cycle number, cycle pair basepair Jaccard index, cycle pair gene Jaccard index, cycle pair overlapping genes (list), cycle pair breakend Jaccard index, cycle pair overlapping breakends (list).

### Analysis module

#### Required file inputs

The **analysis** module requires the following files:

● Cycle level data table with confidence assignments (may be the intra-sample filtered version)
● Jaccard index data table (may be the intra-sample filtered version)
● (optional) Subgroup data table: includes subgroup assignment(s) for each sample in the cycle level data table. Only required if doing subgroup analysis.

### Full cohort visualizations, with single cycle of interest comparisons

The **analysis** module provides visualizations of any structural metrics in the cycle level data table for the full cohort, or a subset of the cohort by confidence assignment. Users can visualize histograms and probability density functions of metrics. They can also compare single cycles of interest to the cohort. We used this section to visualize the cycle size, cycle breakpoint count, and cycle TPR across our full primary BC cohort and high confidence cycles only, highlighting a cycle of interest (**Supplementary figs. 1A-F**).

For example, we inspected a cycle of interest against the primary BC cohort and calculated an empirical p-value to determine the extent to which the cycle deviated. The cycle of interest had a low cycle size, but not significantly lower than the primary BC cohort (p = 0.309, **Supplementary fig. 1A**); it had a breakpoint count above the median of the primary BC cohort, but not significantly greater than the cycle breakpoint counts of the primary BC cohort (p = 0.447, **Supplementary fig. 1B**); it had a high TPR, but not significantly higher than the TPRs of the primary BC cohort (p = 0.164, **Supplementary fig. 1C**). Thus, ecDNAInspector can confirm a cycle has expected characteristics or identify a cycle the user may want to inspect in more detail given outlying characteristics.

This cycle of interest was designated as a high confidence cycle; comparing it against only other high confidence cycles, we clarified its structural features. Calculating the p-value as previously described, this cycle had a more pronounced decreased size relative to the high confidence cycle subset (p_high confidence cycles_ = 0.148, versus p_all cycles_ = 0.309 relative to the full primary BC cohort; **Supplementary fig. 1D**). While it had a higher breakpoint count than the median of the full primary BC cohort, it had fewer breakpoints than the median of the high confidence cycle subset, though the difference was still non-significant (p = 0.486, **Supplementary fig. 1E**). As expected, the cycle did not have significantly higher TPR than the TPRs of the high confidence cycle subset (p = 0.514, **Supplementary fig. 1F**). Thus while the high confidence cycle of interest fit the characteristics of the high confidence cycle subset, its size and breakpoint count trended low, features only clearly visible when comparing to the high confidence cycle subset.

### Pairwise Jaccard index analysis

The ecDNAInspector_analysis module provides a heatmap visualization of all pairwise Jaccard indices (basepair, gene, or breakpoint). It also enables single pair comparison, allowing the user to specify two input cycles and view their pairwise Jaccard indices. We used this section to visualize the basepair Jaccard indices across the full cohort in our primary BC cohort analysis, and compare the Jaccard indices of two high confidence cycle predictions.

### Subgroup analysis

Finally, the ecDNAInspector_analysis module provides visualizations to compare all structural metrics and cycle confidence assignments across user-defined subgroups. We used this section to visualize the differences in structural metrics across the IC subtypes in our primary BC cohort (**Supplementary figs. 5A-H, 7A-F**).

### Validation of ecDNA confidence assignments

#### H3K27Ac HiChIP analysis protocol

The TCGA H3K27Ac HiChIP and ATAC-seq were downloaded from Genomic Data Commons Data Portal. Structural variants (SVs) with breakpoints in regions of copy number gain (CN ≥ 3) were considered potential drivers of aberrant regulatory interactions. Oncogenes were selected based on established cancer gene databases (Bailey et al., *Cell*, 2018). To identify candidate hijacking events, we first intersected ATAC-seq chromatin accessibility peaks with genomic regions showing copy number gains. These peaks were then overlapped with SV breakpoints to pinpoint accessible regions potentially relocated near oncogene promoters due to SVs. Because >75% of promoter–enhancer interactions in the human genome occur within 500 kb (Corces et al., *Science*, 2018), we extended each SV breakpoint by ±500 kb to identify spatially plausible cis-regulatory interactions. We further integrated 1D H3K27ac HiChIP data to validate enhancer activity of these accessible regions. To confirm physical proximity between the hijacked enhancers and oncogene promoters, HiChIP contact maps were analyzed. Only interactions involving known oncogene loci and supported by both regulatory and 3D evidence were retained for downstream analysis.

### Circular structure enrichment and sequencing protocol

The protocol involves extracting high molecular weight (HMW) DNA using HMW DNA extraction kits, followed by Ampure bead purification. Linear DNA is then digested with exonucleases and incubated at 37°C for three days before heat inactivation. Plasmid-Safe ATP-dependent DNAase is added every 24 hours with ATP replenishment. The sample undergoes PEG buffer treatment and magnetic bead separation, followed by incubation with buffer D2 at 65°C and the addition of a STOP solution. Rolling circle amplification (RCA) is performed using REPLI-g sc DNA polymerase at 30°C for 8 hours, followed by polymerase inactivation at 65°C for 3 minutes. The amplified DNA is purified using Ampure beads, eluted in EB buffer, and split into two tubes for Nanopore long-read sequencing and shotgun sequencing. This method enables the capture of both large and small circular DNAs in tumor specimens. DNA input is recorded, and the amplified, exonuclease-digested DNA is quantified before sequencing. The entire procedure takes approximately 8 days from DNA isolation to library preparation. Nanopore sequencing of digested and amplified DNA samples was performed at GeneCore, EMBL Heidelberg, following the manufacturer’s protocol without modifications.

### Predicting cell line ecDNA from short-read sequencing with AmpliconArchitect

We used AmpliconArchitect^13^ (AA, v1.2_r2) to detect ecDNA from short-read sequencing data for the digested UCD12 cell line, digested UCD65 cell line, and non-digested SKBR3 cell line. We used the same specifications as detailed in the “Inferring ecDNA structures from whole genome sequencing” methods section, but aligned to the hg38 assembly instead of the hg19 assembly. This was done to ensure consistency between short-read- and long-read-based cycle predictions, as the tool used for long-read-cycle prediction (CoRAL; see “Predicting cell line ecDNA from long-read sequencing with CoRAL”) requires alignment to the hg38 assembly.

Liftover failed for some cycle segments due to partial sequence deletion in the hg19 assembly; these segments were ignored (note that for our cell line validation, we were interested in consensus genomic regions covered by ecDNA rather than precise segment connections, so the paired breakend prediction impairment resulting from segment deletion was not an issue for our purpose).

<u>Assessing sequencing coverage in predicted short-read ecDNA from linearly-digested samples</u> For each sample, i.e. parental and digested, we binned raw read counts in 50 kB genomic bins across the genome using qDNAseq (v1.26.0). We calculated the average coverage on each bin by multiplying the bin’s read count by 150 (the length of the sequencing reads) and dividing by the bin size (50 kB). We calculated average coverage on the predicted ecDNA segments, extracted from AmpliconArchitect (following the same protocol as detailed in the AmpliconArchitect analysis methods section), for both the digested and parental samples, and normalized this value by their respective median average coverage across all bins. We generated null “ecDNA regions” (i.e., simulated ecDNA regions composed of randomly generated genomic regions of the same size and number as the original ecDNA) only from mappable regions on the genome, using http://hgdownload.soe.ucsc.edu/gbdb/hg19/bbi/problematic/ to remove unmappable regions. To do so, we randomly selected a chromosome followed by a genomic position within that chromosome’s mappable regions. If a segment of the desired length could be obtained from this starting position, we stored it as a segment of the null ecDNA. If the mappable region ended before the full length of the predicted ecDNA segment could be matched, we randomly sampled a new start position. Once all length-matched segments were generated, we returned the null “ecDNA regions”. We ran the same procedure for 1000 iterations. For each sample, we compared the average coverage in the predicted ecDNA region to this null distribution and calculated a p-value by subtracting the proportion of null log ratios that were less than the predicted log ratio from 1. See **Supplementary fig. 6B**.

### Predicting cell line ecDNA from long-read sequencing with CoRAL

We used CoRAL^15^ (v1.0.0) to generate cycle predictions for the UCD12 and UCD65 linearly digested cell lines. First, we used seqfilter (https://github.com/clwgg/seqfilter) to filter all fastq files with fewer than 10,000 reads using the following parameters: -n -m 10000 -i in.fq -o out.fq. Next, we used nextflow (v22.10.5) to run the nf-core/nanoseq pipeline to align the fastq files using the following parameters: --input samplesheet.csv --genome GRCh38 -profile singularity - c config_file -r 3.1.0 --protocol DNA --skip_basecalling --skip_demultiplexing --skip_vc --skip_sv --skip_quantification --skip_differential_analysis. This was run with java 17.0.6.

Resulting bam files for each cell line were merged using samtools (v1.21). Next, candidate amplifications were detected using CNVKit (v0.9.10) with the default circular binary segmentation (CBS) setting. The resulting copy number segment file was used to seed CoRAL with minimum breakpoint support set to 2.0 (--min_bp_support 2.0).

### Cell line validation of cycle confidence assignments

For short-read- and long-read-based cycle predictions for the UCD12 and UCD65 cell lines and short-read-based cycle predictions for the SKBR3, we performed a liftover (https://genome.ucsc.edu/cgi-bin/hgLiftOver) from the hg38 assembly to the hg19 assembly to match our primary BC cohort cycle prediction coordinates. For both the UCD12 and UCD65 cell lines, we subsetted all short-read- and long-read-based cycles to those containing the IC6 cytoband chromosome (chromosome 8) and with size greater than 1 Mbp. We computed the “consensus” ecDNA regions for each cell line by finding the intersection of all short- and long-read-based ecDNA regions for that cell line (**Figure 3A, Supplementary fig. 6A**). These regions were saved as the short-read long-read consensus ecDNA regions. The basepair Jaccard index between these consensus regions and each unique IC6 cycle was computed and stored for each cell line. We repeated this analysis using the ecDNA cycles predicted from the SKBR3 cell line short-read sequencing data, computing the basepair Jaccard index between each SKBR3 cycle and each IC6 cycle (**Figure 3D-E, Supplementary figs. 6C-D**).

### Application of ecDNAInspector to IC-subtyped Breast Cancer Cohort

Application: cycle metric calculation, cycle clustering, cycle confidence assignments, and intrasample filtering modules with ecDNAInspector_cycle_selection notebook

To analyze our IC-subtyped Breast Cancer Cohort, we started by using the **ecDNAInspector_cycle_selection** module in a Jupyter notebook. We input file and directory paths to all our AA-output cycle prediction files, our orthogonal SV calls bedpe file, our gene information bed file (we used the hg19 v98 Cosmic Cancer Gene Census [https://cancer.sanger.ac.uk/cosmic/download/cosmic/v98/cancergenecensus]), and our genome “blacklist” information bed file (we used the hg19 Genome Blacklist [https://github.com/Boyle-Lab/Blacklist/tree/master/lists]). We also specified the output file paths for the cycle level data table (includes all structural metric information), the cycle level data table with clustering (includes all structural metric information and the results of the clustering), the cycle level data table with confidence (includes all structural metric information, the results of the clustering, and the final cycle confidence assignments [determined by cluster or by the optional additional filtering step]), and the intra-sample filtered cycle level data table with confidence (the subset of the full cycle level data table with confidence, excluding lower quality cycle “copies” [see “Optional intra-sample similarity filtering”]). Because we were using AA-output cycle predictions, we specified to include “all” cycle types (rather than “circular” or “linear” only; see “General structural feature extraction”). Finally, we specified all buffers, thresholds, and parameters as follows:

● be_overlap_buffer = 100; maximum distance between breakends to be overlapping
● SV_in_range_buffer = 100; maximum distance between structural variant and breakend to be overlapping
● blacklist_buffer = 100; maximum distance between breakend and blacklist region for breakend to be considered included
● small_del_len = 2000; maximum length of deletion to be excluded from SV validations
● gene_inclusion_prop_buffer = 1.0; proportion of total gene length that must be present in cycle for full gene to be considered included in cycle
● copy_count_threshold = 4.0; minimum copy count to include single cycle
● duplicate_buffer = 100; maximum distance between matched ends of paired ends for paired ends to be considered duplicates
● min_clusters = 3; minimum number of clusters to try in consensus clustering
● max_clusters = 5; maximum number of clusters to try in consensus clustering
● cluster_num_resamples = 1000; number of resamples for consensus clustering iterations
● cluster_resample_prop = 0.8; proportion of samples to resample for consensus clustering iterations
● n_init = 10; number of KMeans runs with different centroid seeds

Following the clustering, we elected to use the optional intra-sample filtering step with a similarity threshold = 0.2 (see “Optional intra-sample similarity filtering”).

<u>Application: Jaccard calculation module with ecDNAInspector_Jaccard_calculation notebook</u> We then used the **ecDNAInspector_Jaccard_calculation** module to calculate the pairwise Jaccard indices (basepair, gene, and breakend) between cycles across different samples. We input the path to the intra-sample filtered cycle level data table from the previous module, the path to an output file to store all Jaccard information, and defined the breakend overlap buffer = 100 base pairs (i.e., breakends on different cycles within 100 basepairs of each other would be considered overlapping).

### Application: Analysis module with ecDNAInspector_analysis notebook

Finally, we used the **ecDNAInspector_analysis** module to perform cohort-level visualizations of structural metrics and Jaccard indices, single-cycle versus cohort comparisons, and subgroup comparisons across the IC subtypes (see “ecDNAInspector_analysis module”). We input paths to the intra-sample filtered cycle level data table with confidence information, the intra-sample filtered Jaccard index data table, and a table with subgroup information (i.e., the IC subtype) for each sample.

### Data calculations, visualizations, and statistics

For all data calculations, visualizations, and statistics, ecDNAInspector uses the following packages: numpy (v1.24.0), pandas (v2.2.2), matplotlib (v3.9.2), seaborn (v0.12.2), scipy (v1.15.2). ecDNAInspector was run using Python 3.10.9 for our primary BC cohort analysis.

## Data Availability

All cohorts are publicly available. Data for TCGA BRCA samples can be found on the Genomic Data Commons Data Portal (https://portal.gdc.cancer.gov/). DNA-sequencing data for the International Cancer Genome Consortium (ICGC) breast cancer samples can be found on the European Genome-Phenome Archive (accession numbers EGAD00001000141, EGAD00001001322, EGAD00001001334, EGAD00001001335, EGAD00001001336, EGAD00001001337 and EGAD00001001338). In cases in which it was possible, alignments for both TCGA and ICGC samples carried out by the Pancancer Analysis of Whole Genomes were used (https://docs.icgc-argo.org/docs/data-access/icgc-25k-data).

## Code Availability

The ecDNAInspector tool is available through GitHub for open use at https://github.com/cancersysbio/ecDNAInspector. In addition, code for computational analysis would be made available via the Curtis Lab GitHub repository upon publication.

