## Supplemental Figures for "High-confidence structural predictions of extrachromosomal DNA with ecDNAInspector"

### Supplementary Figures

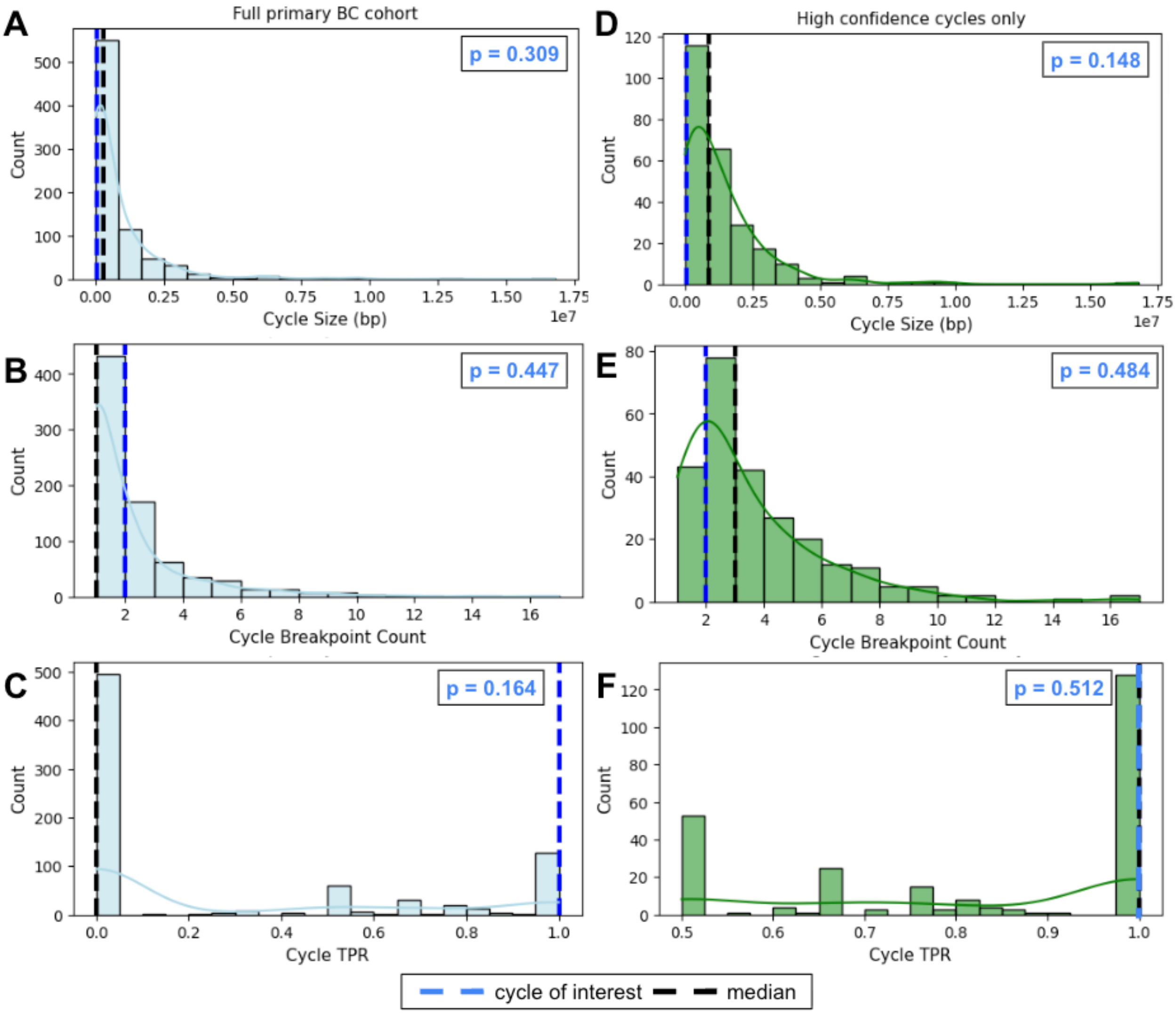


**Supplementary Figure 1. ecDNAInspector highlights cycle of interest against full primary BC cohort and high confidence cycle subset**. The ecDNAInspector_analysis module visualizes metric distributions as count histograms and probability density functions (PDFs) for the full primary BC cohort (left column) and high confidence cycle subset (right column). Metrics for a single cycle of interest (highlighted in blue) are compared to these distributions. Metrics include: A) cycle size (full primary BC cohort), B) cycle breakpoint count (full primary BC cohort), C) cycle TPR (full primary BC cohort), D) cycle size (high confidence cycle subset), E) cycle breakpoint count (high confidence cycle subset), and F) cycle TPR (high confidence cycle subset). P-values are calculated by determining the proportion of cohort metric values that were less than (if the cycle metric trended to the right of the cohort distribution) or greater than (if the cycle metric trended to the left of the cohort distribution) the cycle metric, and subtracting this proportion from 1.

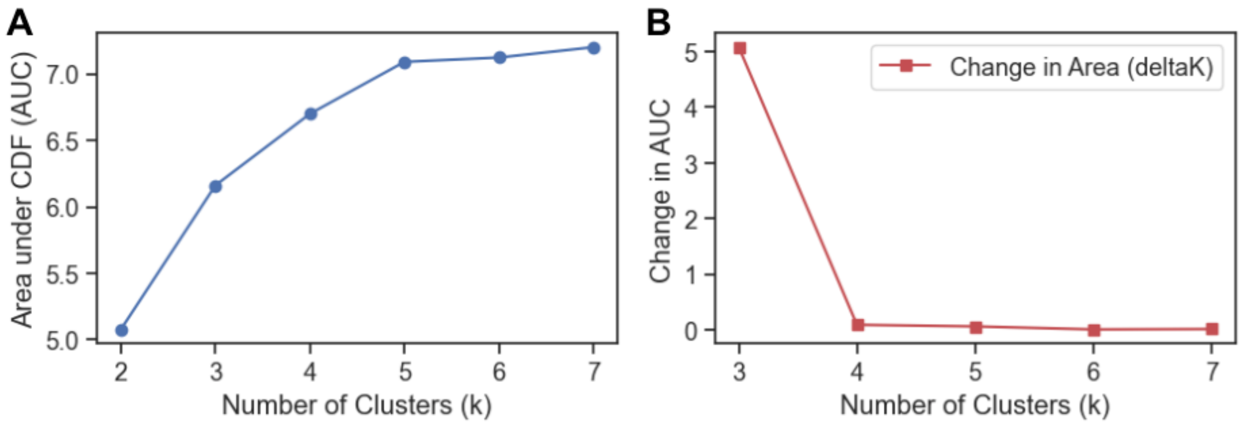


**Supplementary Figure 2. Consensus clustering finds K = 3 as optimal number of clusters for primary BC cohort**. Plots of area under the CDF (AUC) (A) and the change in AUC (B) find K = 3 as a stable number of clusters, with additional clusters resulting in minimal gains in clustering stability.

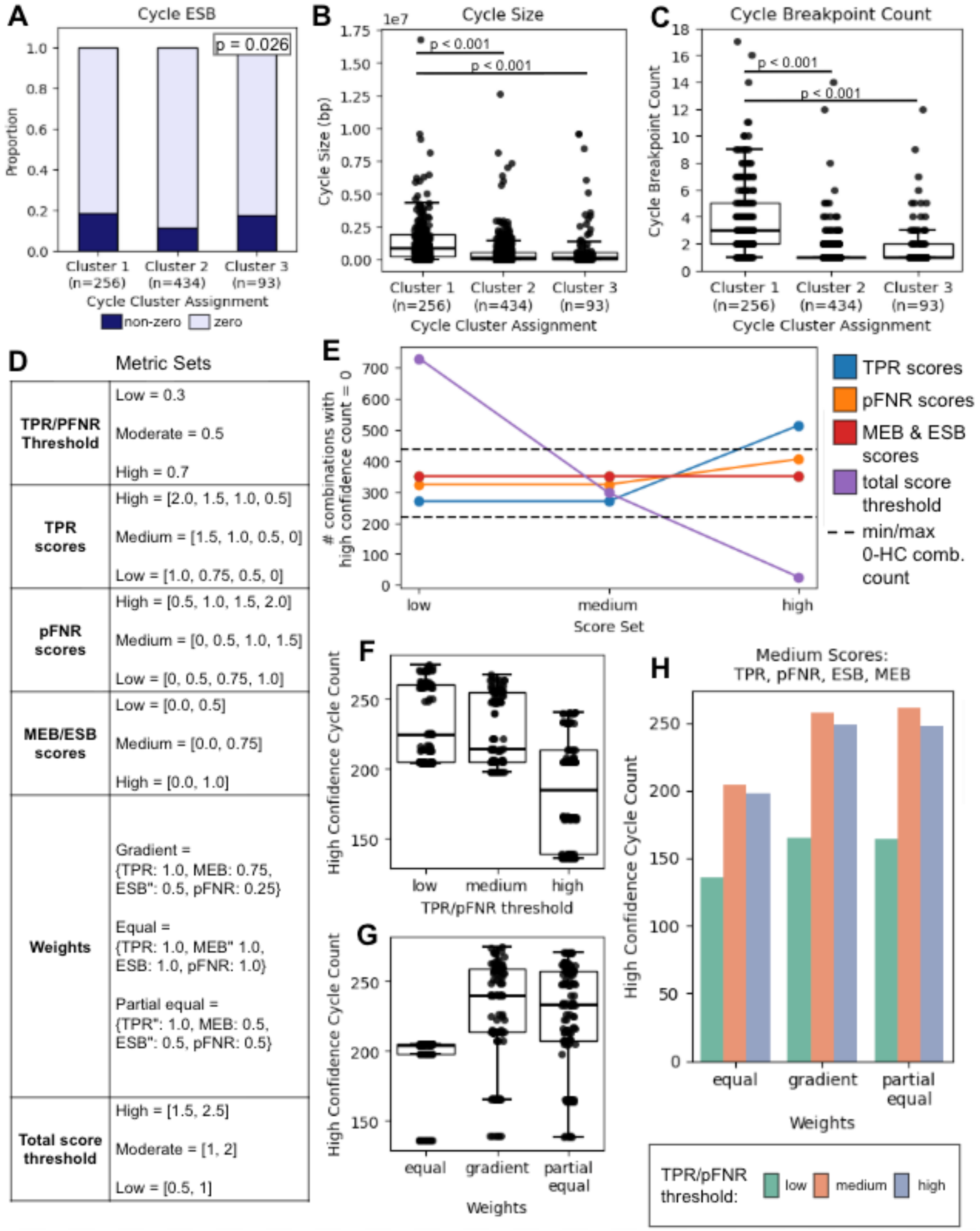


**Supplementary Figure 3. ecDNAInspector clustering and filtering validated by cluster cycle types and filtering output stability.** Plots show the difference in A) cycle ESB, B) cycle size, and C) cycle breakpoint count. P-values calculated with Chi-squared test (A) and Kruskal-Wallis test with post-hoc Dunn test (B, C). D) Different combinations of metric sets were tested for ecDNAInspector's optional filtering step. E) The count of metric set combinations producing 0 high confidence cycle assignments is shown for different metrics. Each dot indicates the count of metric set combinations producing 0 high confidence cycle assignments while holding one metric set constant at the x-axis value. For each metric, we continued to test only metric set values that produced a count within 10-20% of the total combinations (range denoted by dashed horizontal lines). This eliminated the use of low and high total score thresholds, and high TPR scores. We also eliminated the use of high pFNR scores. F) The distribution of the count of high confidence cycles (y-axis) produced with different combinations of remaining metric sets while holding the TPR/pFNR thresholds constant at the x-axis value. G) The distribution of the count of high confidence cycles (y-axis) produced with different combinations of remaining metric sets while holding the weights constant at the x-axis value. I). The count of high confidence cycles produced while varying the weights and TPR/pFNR threshold, holding the TPR, pFNR, ESB, and MEB scores constant at medium. Medium TPR/pFNR/ESB/MEB scores, medium TPR/pFNR threshold, and gradient scores were selected to filter our IC-subtyped Breast Cohort. Boxplots represent median, 0.25 and 0.75 quantiles with whiskers at 1.5x the interquartile range.

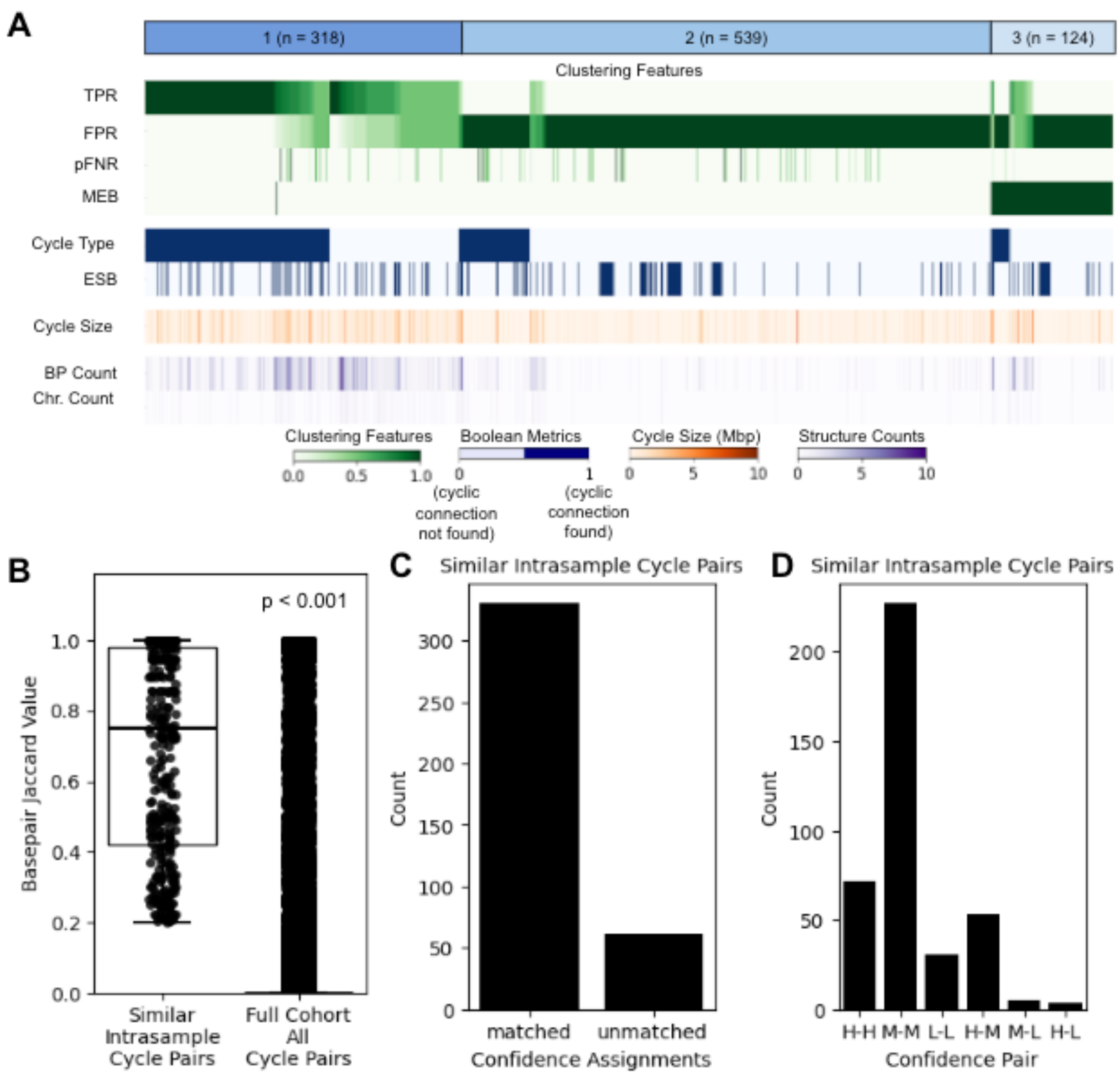


**Supplementary Figure 4. Intra-sample filtering reduces same-sample cycle redundancy without altering clustering or confidence assignments.** A) Heatmap of metrics by cluster for ecDNAInspector clustering on IC-subtyped Breast Cohort, without intra-sample filtering step. Shows distinguishing structural features by cluster. B) Basepair Jaccard indices for similar intrasample cycle pairs (i.e., cycle pairs from the same sample with BP Jaccard index > 0.2) versus basepair Jaccard indices for all inter- and intra-sample cycle pairs. P-value calculated with two-sided Mann-Whitney U-test. C, D) Shows count of cycle pairs from the same sample with >20% similarity that receive the same confidence assignment (matched) or different confidence assignments (unmatched). Matches are split by confidence set in D), displaying counts for high-high (H-H) match, medium-medium (M-M) match, low-low (L-L) match, high-medium (H-M) mismatch, medium-low (M-L) mismatch, and high-low (H-L) mismatch.

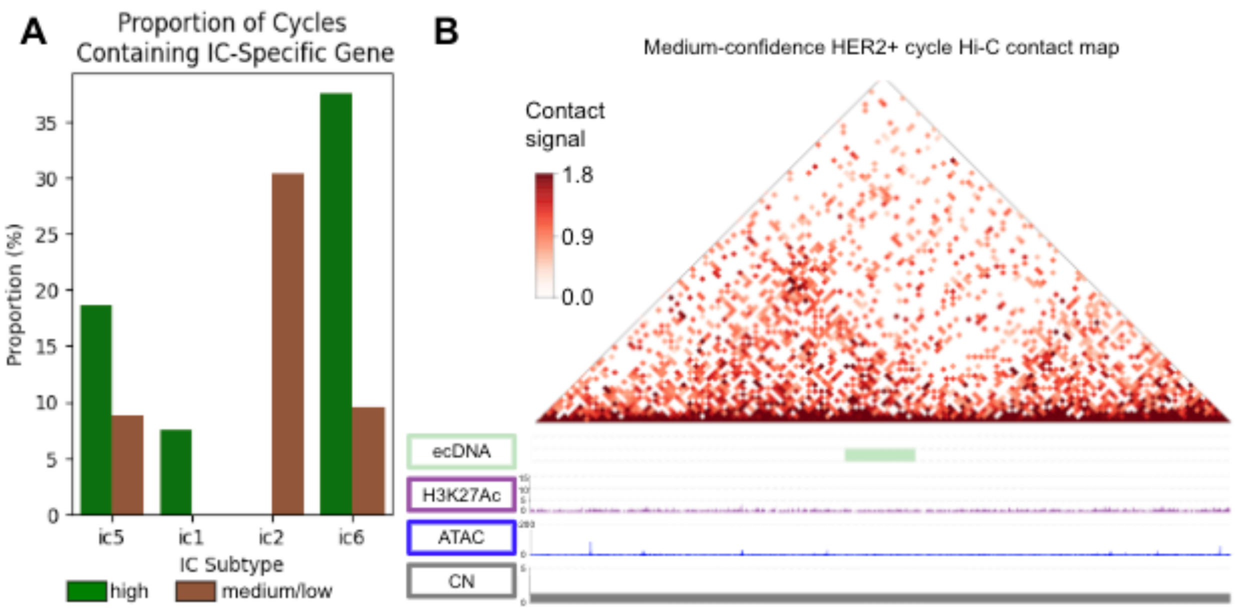


**Supplementary Figure 5. ecDNAInspector confidence assignments validated by gene inclusion and structural contacts.** A) IC-specific gene inclusion in cycle predictions from IC-subtype matched patients is shown by cycle prediction confidence. B) H3K27ac and ATAC HiChIP contact matrix (10-kb resolution) in a medium confidence cycle from a HER2+ patient, with only sparse structural proximity of cycle regions to each other. Copy number (CN) is ploidy-corrected.

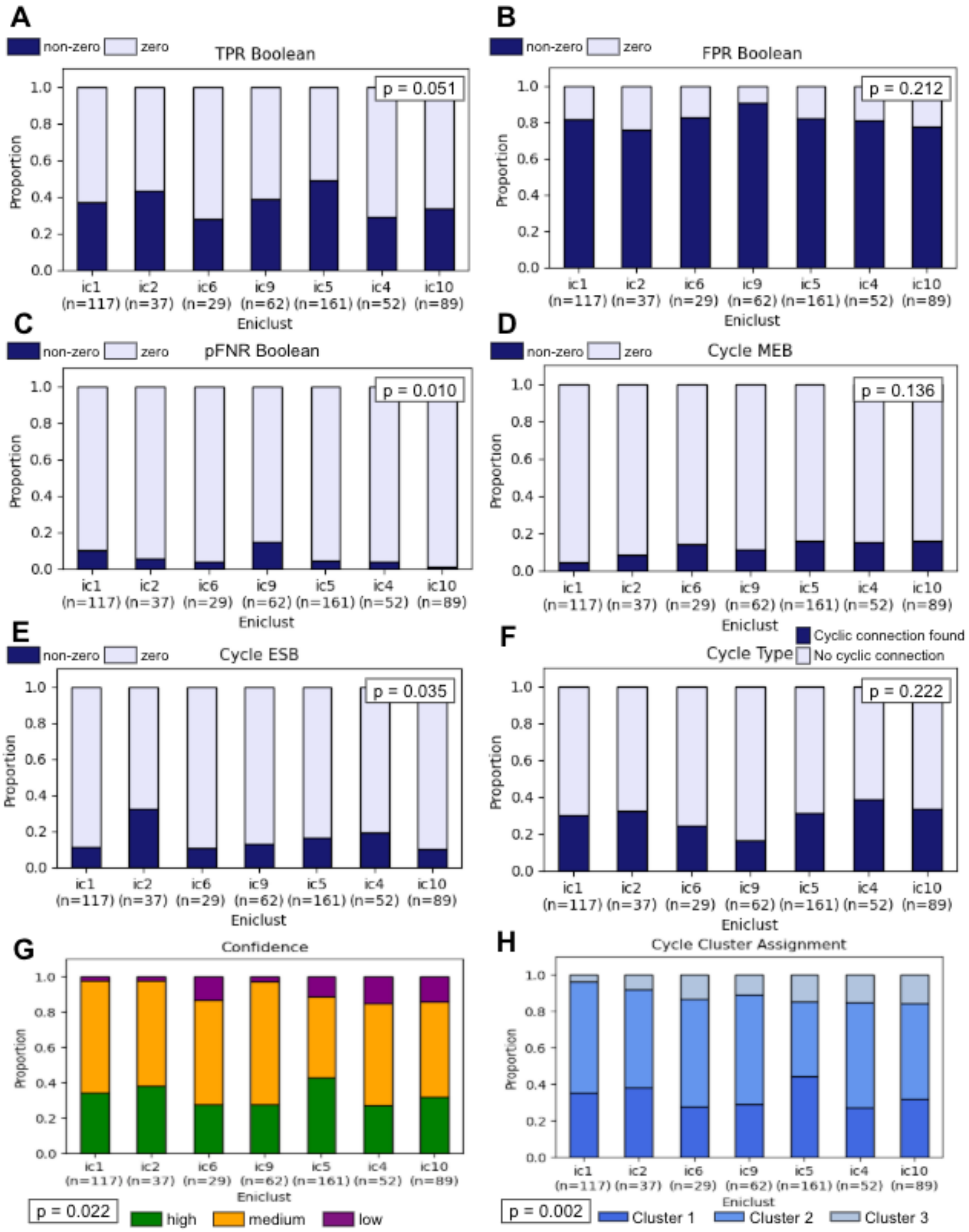


**Supplementary Figure 6. Structural metrics, clustering, and confidence assignments stable across IC subtypes.** Plots of A) TPR Boolean, B) FPR Boolean, C) pFNR Boolean, D) Cycle

MEB, E) Cycle ESB, F) Cycle Type, G) Cycle Confidence, and H) Cycle Cluster Assignment show differences in ecDNAInspector-calculated metrics and clustering/filtering output between IC subtypes. P-values calculated with Chi-squared tests.

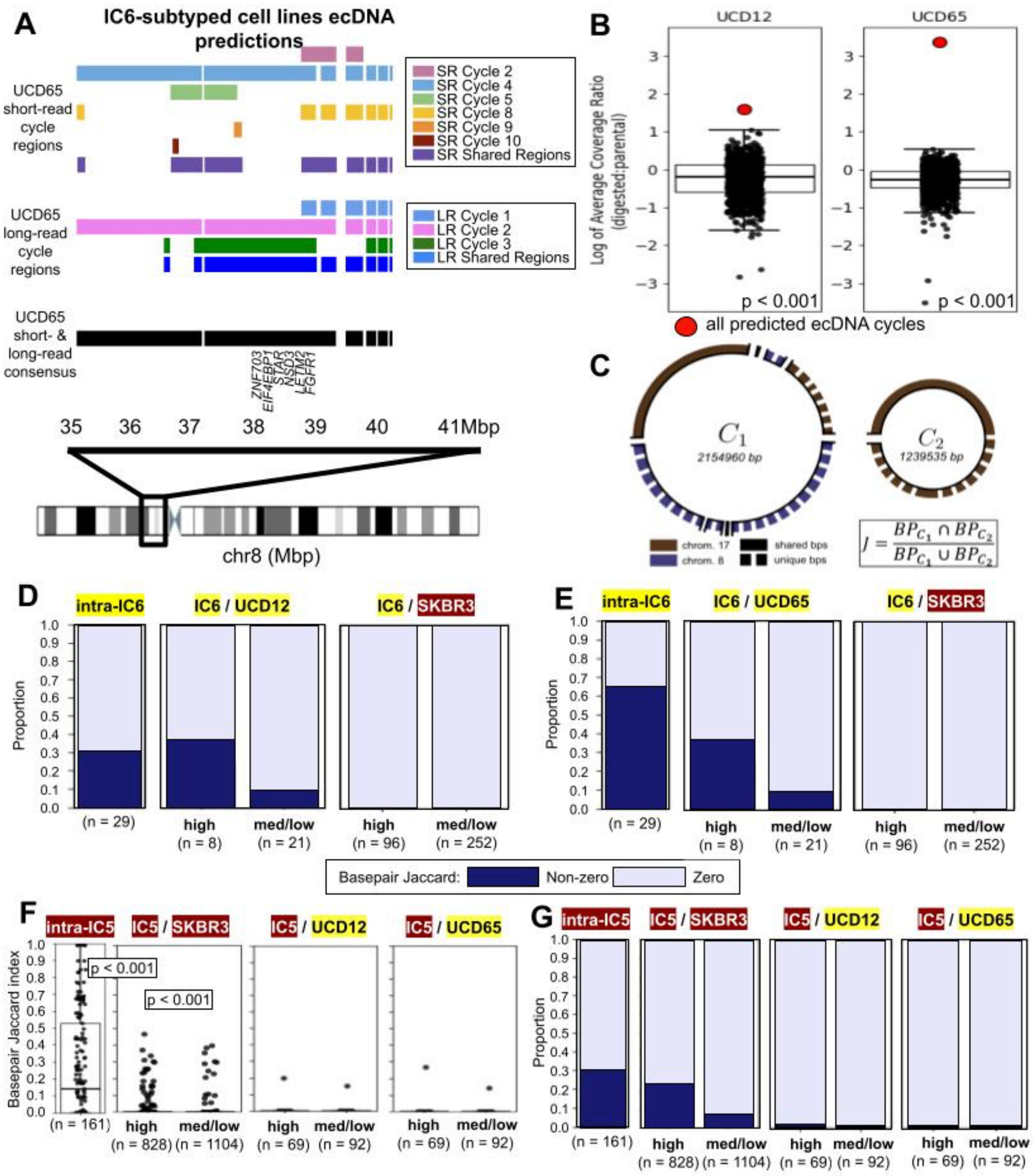


**Supplementary Figure 7. ecDNA-enriched cell lines validate IC-subtyped Breast Cohort cycle confidence assignments.** A) A schematic of genomic regions in ecDNA cycle predictions based on short-read and long-read sequencing of the UCD65 cell line. Notable oncogenes are specified. Short-read and long-read regions are color-coded by whether the cycle number they are found in, or whether regions are shared across cycles. B) Ratio of sequencing coverage in digested versus parental UCD12 and UCD65 cell lines in the predicted ecDNA regions (red dot)

compared to 1,000 null regions. Boxplots represent median, 0.25 and 0.75 quantiles with whiskers at 1.5x the interquartile range. P-value calculated as the proportion of null log ratios that were less than the predicted ecDNA log ratio, subtracted from 1. C) A schematic detailing how pairwise basepair Jaccard index metric is calculated. D, E) Proportions of non-zero or zero basepair Jaccard indices for intra-IC6 cycle comparisons, IC6-UCD12 (C) or UC6-UCD65 (D) cycle comparisons split by confidence, and IC6-SKBR3 cycle comparisons split by confidence. F) The basepair Jaccard index is used to assess similarity of IC5 high confidence and grouped medium and low confidence cycle predictions to ecDNA regions from the IC5-subtyped SKBR3 cell line and IC6-subtyped UCD12 and UCD65 cell lines. G) Proportions of non-zero or zero basepair Jaccard indices for intra-IC5 cycle comparisons, IC5-SKBR3 cycle comparisons split by confidence, and IC5-UCD12 and IC5-UCD65 cycle comparisons split by confidence. P-values calculated with Mann-Whitney U-test. Boxplots represent median, 0.25 and 0.75 quantiles with whiskers at 1.5x the interquartile range.

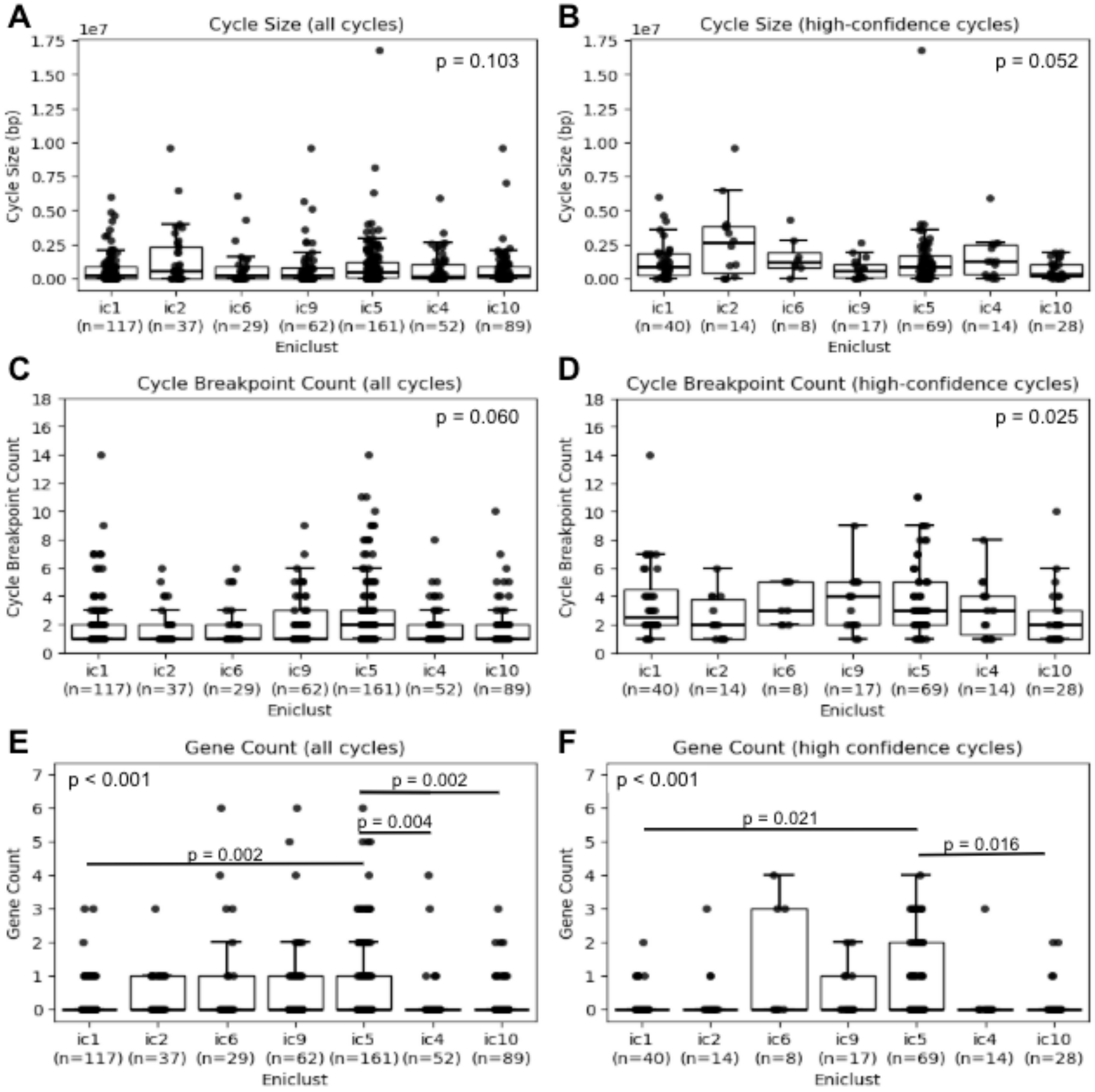


**Supplementary Figure 8. Subsetting analysis to high confidence cycles clarifies structural metrics differences across full cohort and cohort subgroups**. Cycle size was not significantly different across IC subtypes, both when considering all cycles (A) and subsetting to high confidence cycles only (B). Cycle breakpoint count was not significantly different across IC subtypes, both when considering all cycles (C) and subsetting to high confidence cycles only (D). Gene counts in all cycles were significantly different across IC subtypes (E). This pattern was confirmed in high confidence cycles only (F). All p-values calculated with Kruskall-Wallis tests and post-hoc Dunn tests (E, F). Boxplots represent median, 0.25 and 0.75 quantiles with whiskers at 1.5x the interquartile range.

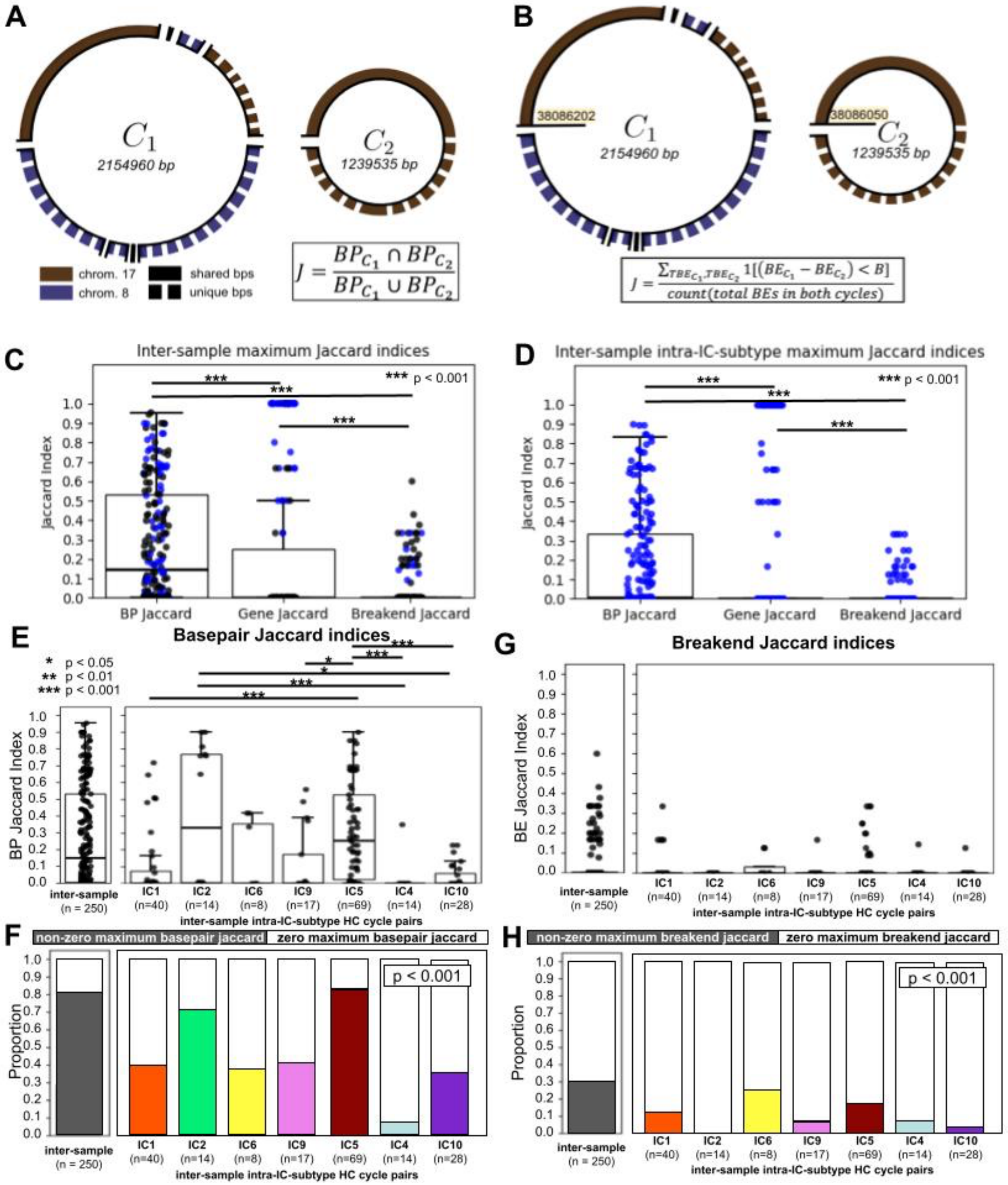


**Supplementary Figure 9. ecDNAInspector analysis module shows strong basepair conservation and weak breakend conservation across IC subtypes**. A) A schematic detailing how pairwise basepair Jaccard index metric is calculated. B) A schematic detailing how pairwise breakend Jaccard index metric is calculated. Basepair, gene, and breakend Jaccard indices were calculated for every cycle pair across samples across samples. The maximum Jaccard index achieved by each high confidence cycle was plotted (C), with basepair Jaccard indices being significantly greater than both gene Jaccard indices and breakend Jaccard indices. Gene Jaccard indices were also significantly greater than breakend Jaccard indices. Jaccard indices for cycle pairs from the same subgroup (intra-subgroup cycle pairs) were plotted (D), with trends following those seen in C. Blue dots indicate maximum Jaccard values resulting from intra-IC-subtype cycle pairs. Cycle pairs were split across subgroups. E) Inter-subgroup differences in basepair Jaccard indices were significant ($p < 0.001$, Kruskal-Wallis test), with several pairwise groups showing significant differences. F) Proportions of non-zero versus zero maximum basepair Jaccard indices for inter-sample cycle pairs and intra-subgroup inter-sample cycle pairs were also compared. Inter-subgroup differences were significant. G, H) Inter-subgroup differences in breakend Jaccard indices were not significant. P-values calculated with pairwise two-sided Mann-Whitney U-tests (C, D), Kruskall-Wallis tests (E, G) and post-hoc Dunn tests (E), and Chi-squared tests (F, H). Boxplots represent median, 0.25 and 0.75 quantiles with whiskers at 1.5x the interquartile range.
